# A double-assurance mechanism restrains generation of high potential transit-amplifying progenitors during neurogenesis

**DOI:** 10.1101/2025.09.16.676671

**Authors:** Cyrina M. Ostgaard, Arjun Rajan, Sophia R. Davidson, Cheng-Yu Lee

## Abstract

Stem cells can generate differentiated progeny directly or indirectly through transit-amplifying progenitors (TAPs), which are vulnerable to tumorigenic transformation. Despite clinical relevance, how stem cells regulate TAP production is unclear. *Drosophila* brains contain Asense^+^ stem cells (Type1 neuroblasts; T1NBs) that generate low neurogenic potential TAPs and Asense^-^ stem cells (Type2 neuroblasts; T2NBs) that produce high-potential TAPs [Intermediate Neural Progenitors (INPs)]. Unexpectedly, cell-type-specific enhancers of genes essential for INP formation are poised in T1NBs despite never generating INP progeny. Thus, both NB types are competent to generate INPs. Inducing T1NBs to generate INPs is more efficient upon *asense* knockdown. The progeny of T2NBs expressing elevated Asense adopt a low-potential TAP identity instead of INPs. Elevated Asense expression limits tumor NB expansion driven by stalled INPs halting tumor expansion. We propose that a double-assurance mechanism characterized by dynamic superimposition of lineage-specific activators upon basal-state determinants defines NB capability to generate high-potential TAPs.

## Introduction

TAPs are transient stem cell progeny that function to increase the output of differentiated cells during development and homeostasis. TAPs can vary widely in terms of proliferative and developmental potential.^1^ During vertebrate neurogenesis, radial glial progenitors function as neural stem cells and can generate neurons indirectly through TAPs, also known as intermediate progenitors.^2–6^ Most intermediate progenitors display low neurogenic potential and canonically divide once to produce two neurons. In developing primate and human brains, a subset of intermediate progenitors exhibit higher neurogenic potential and can divide multiple times to increase neuron generation.^7–9^ It remains unclear whether chromatin accessibility or histone modification patterns at enhancers of genes that drive the formation of specific intermediate progenitors define radial glial capability to generate high- or low-potential TAP subtypes.

The fly larval brain contains two distinct NB lineages that are well-defined providing an excellent *in vivo* paradigm for investigating how heterogenous stem cells regulate generation of distinct types of TAPs.^10–12^ Approximately 100 NBs are specified per brain lobe during embryogenesis and undergo repeated asymmetric division to self-renew and to produce an identical TAP subtype every division.^13,14^ Most of these are T1NBs, which generate ganglion mother cells (GMCs) that exhibit low neurogenic potential and divide only once to produce two neurons. Eight are T2NBs, which invariably produce an immature INP that is Asense (Ase; Ascl1) negative but will upregulate Ase as the differentiation to an INP progresses.^15–17^ An INP exhibits high neurogenic potential compared to a GMC. An INP undergoes 5-6 rounds of asymmetric division, generating a GMC each division, leading to an overall output of more than a dozen differentiated cells including neurons and glia in its lifetime.^18,19^ Due to the invariability of the NB progeny and that which NBs produce INPs is determined when the NBs are first generated during the embryonic stage, the ability to generate INPs is thought to be genetically determined by the developmental program that specifies T2NB identity. This “deterministic model” predicts that nucleosomes at enhancers that drive the expression of genes essential for INP formation are closely spaced from each other (chromatin inaccessible) making these *cis*-regulatory elements inaccessible to TF binding in T1NBs resulting in the lack of INP progeny.

Post-embryonically removing *trithorax* (*trx*) gene activity converts T2NBs to T1NBs as indicated by (1) loss of T2NB-specific marker expression, (2) gain of T1NB-specific gene expression and (3) direct generation GMCs.^20^ Trx is an evolutionarily conserved histone methyltransferase and maintains chromatin in an active state through H3K4 methylation at target gene loci.^21,22^ Trx maintains expression of transcription factors (TFs) essential for T2NB identity. These genes include Tailless (Tll) which is necessary and sufficient for T2NB identity with high levels of Tll expression promotingT2NB maintenance, likely by suppressing the T1NB genetic program.^23,24^ Pointed (Pnt) promotes immature INP differentiation into INPs.^20,25–27^ Additionally, T2NBs mutant for *buttonhead* (*btd*) generate GMCs instead of INPs while maintaining T2NB-specific marker gene expression.^20,26^ *btd* encodes a C_2_H_2_ zinc finger TF that likely functions as a positive regulator of gene expression.^28–30^ Mis-expressing *btd* is sufficient to induce T1NBs to generate INPs. These data suggest that GMC identity is the basal-state for NB progeny fate in both lineages. This “hierarchical model” predicts that chromatin at enhancers that drive gene expression essential for INP formation are loosely wrapped around histone octamers (chromatin accessible) but post-translational modifications of histones maintain inactivity of these *cis*-regulatory elements (chromatin state) in T1NBs. Trx, Tll and Btd can promote activation of these cell-type-specific enhancers in T2NBs leading to generation of INPs instead of GMCs.

The basic-helix-loop-helix TF Ase is expressed in T1NBs throughout development and has been implicated in determining T1NB lineage identity.^13–17,31–33^ Surprisingly, deletion of the *ase* locus has no effect on viability or on T1NB lineage progression. This suggests that Ase is not necessary to maintain T1NB identity or for the generation of GMCs. Although increasing Ase expression is sufficient to block premalignant T2NB formation when Tll is overexpressed in T1NBs, the functional significance and underlying mechanism of this result is currently unclear.^23^ As such, Ase mainly serves as a reliable marker for identifying T1NBs and INPs.

We hypothesized that the chromatin state at enhancers that drive gene expression in immature INPs to promote INP formation serves as a functional readout for the capability of NBs to generate INPs. Thus, identifying genes that are highly transcribed in immature INPs and defining enhancers that drive their cell-type-specific expression are critical for investigation of NB capability to generate INPs. We leveraged published single-cell mRNA sequencing datasets of wild-type NB lineages ^34–36^ to identify genes that are high transcribed in immature INPs. We searched for active neurogenic enhancers, which are defined as non-promoter *cis*-regulatory regions that display accessible chromatin and are bound by Zelda (Zld; a positive regulator of gene transcription ^37,38^) and Fruitless^C^ (Fru^C^; a negative regulator of gene transcription ^36^) in T2NBs. We identified 3839 active neurogenic enhancers including cell-type-specific enhancers associated with genes that promote immature INP differentiation to INPs. Enhancers of genes essential for INP formation and functions display features that are typically associated with poised enhancers including accessible chromatin, high H3K27me3/H3K4me1 and low H3K27Ac in T1NBs. Thus, the chromatin state and not accessibility at enhancers associated with genes essential for INP formation and functions correlates with NB capability to give rise to an INP or GMC. Inducing T1NBs to generate INPs by mis-expressing Btd becomes more efficient when *ase* function is knocked down, while high levels of Ase promote T2NBs to generate GMCs, via bypassing the INP stage. We identified the TF Prospero (Pros) as the key Ase downstream-effector that reprograms T2NBs to generate GMCs by activating the GMC genetic program in immature INPs. Furthermore, elevating Ase expression in tumor NBs that form a continual positive feedback loop with immature INPs that are stalled in differentiation suppresses tumor expansion. These results led us to propose a double-assurance mechanism in which dynamic superimposition of lineage-specific activators (Btd) upon basal-state determinants (Ase and Pros) defines NB capability to generate INPs or GMCs.

## RESULTS

### Transcriptional states regulate NB capability to generate INPs

To determine whether chromatin accessibility or post-translational modification of histones at enhancers of genes essential for INP formation correlates with the capability of NBs to generate INPs (Fig. 1A), we first defined gene transcripts that are high enriched in T2NBs, immature INPs or INPs. Of the 255 genes which showed transcript enrichment in T2NBs, immature INPs or INPs based on previously published single-cell mRNA sequencing datasets of wild-type larval brains ^34–36^, 174 showed region of open accessibility (Table S1). These genes are assigned to one of four groups based on the cell type where their transcripts first become detectable (Fig. 1B; Table S1). Transcripts encoded by the Group 1 genes (e.g. *btd* and *pnt*) become detectable prior to or coinciding with the onset of immature INP differentiation marked by Earmuff (Erm) protein expression.^20,26^ Gene transcripts in the Group 2 (e.g. *erm*) were first detected in the early stage of immature INP differentiation (Erm^+^Ase^-^).^27,39,40^ Gene transcripts in the Group 3 (e.g. *hamlet* (*ham*), *Dichaete* (*D*)) are detected starting in the late stages of immature INP differentiation (Erm^+^Ase^+^).^18,24^ Transcripts encoded by the Group 4 genes (e.g. *odd-paired* (*opa*) and *eyeless* (*ey*)) are detected in various INP life stages.^18,41^ Combined, there are 81 genes in group 3 & 4 that are highly enriched during immature INP differentiation and 62 have associated chromatin accessible regions. Assessing the chromatin state at enhancers that drive their cell-type-specific expression should unravel whether chromatin accessibility or histone modifications correlates with NB capability to generate INPs.

**Figure 1.**
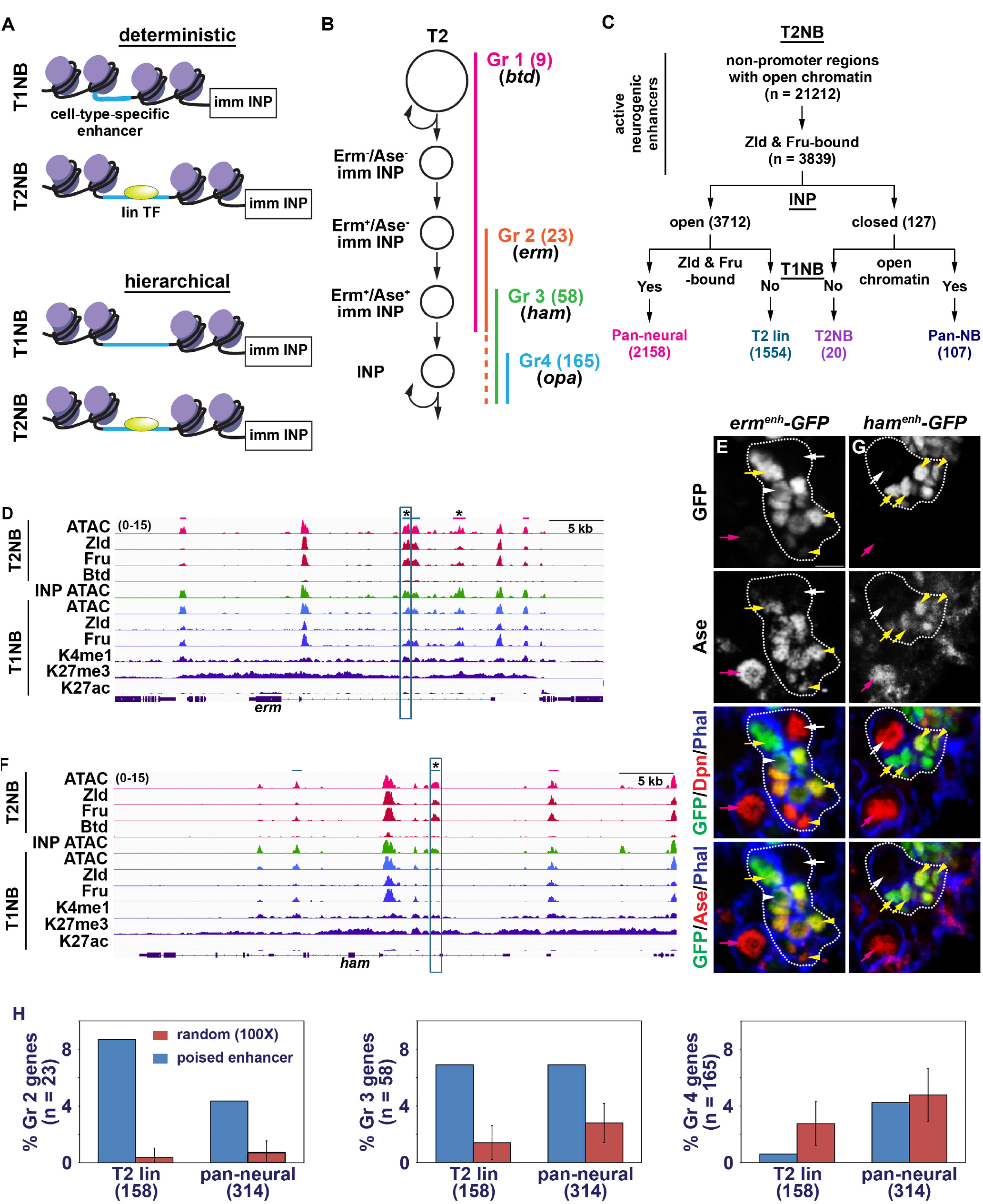
Cell-type-specific enhancers associated with genes essential for INP formation are poised in T1NBs. (A) Models for the regulation of NB capability to generate INPs. The deterministic model holds true if cell-type-specific enhancers associated with genes that function to promote immature INP (imm INP) differentiation to are inaccessible in T1NBs. The hierarchal model holds true if imm INP enhancers are accessible but inactive in T1NBs. Lineage-specific transcription factor (lin. TF) promotes activation of these enhancers in T2NBs. (B) 255 genes showed transcript enrichment in T2NBs, immature INPs or INPs based on previously published single-cell mRNA sequencing datasets of wild-type larval brains.^34–36^ These genes are assigned to one of four groups based on the cell type in which their transcripts first become detectable. (C) Flowchart for the identification of cell-type-specific enhancers in fly larval brain NB lineages. Pan-neural: Pan-neural enhancer. T2 lin: T2 lineage enhancer. T2NB: T2NB enhancer. Pan-NB: pan-NB enhancer (D,F) Genome browser track of the *erm* and *ham* locus showing patterns of chromatin accessibility, TF occupancy and histone marks in indicated cell types. The magenta bars correspond to pan-neural enhancers, and teal bars correspond to T2-lin enhancers. Teal box designates enhancer that recapitulates the onset of *erm* or *ham* expression in the T2 lineage. * indicates poised enhancers. (E) GFP reporter expression driven by an *erm* T2 lin enhancer (teal box in D) becomes detectable in Ase^-^ immature INPs but is undetectable in T1 and T2NBs. (G) GFP reporter expression driven by a *ham* T2 lin enhancer (teal box in F) becomes detectable in Ase^+^ immature INPs but is undetectable in T1 and T2NBs. White dotted line circles T2NB lineage. White arrow: T2NB. White arrowhead: Ase^-^ immature INP. Yellow arrow: Ase^+^ immature INP. Yellow arrowhead: INP. Magenta arrow: T1NB. Scale bar: 10µm.

We took an unbiased approach to identify active neurogenic enhancers, which are defined as non-promoter *cis*-regulatory regions that display accessible chromatin and are bound by Zld and Fru^C^ in T2NBs using published datasets.^36,38^ 3839 *cis*-regulatory regions fulfilled these criteria and were referred to as active neurogenic enhancers in T2NBs (Fig. 1C). Chromatin accessibility in INPs separates these enhancers into the NB-specific cohort from the NB lineage cohort. Enhancers in the NB-specific cohort are divided into Pan-NB enhancers or T2NB enhancers based on their chromatin accessibility in T1NBs (Fig. 1C). Pan-NB enhancers display accessible chromatin in T1NBs and likely promote gene expression in both NB types (Fig. S1B; Table S2). T2NB enhancers display inaccessible chromatin in T1NBs and likely regulate the expression of genes (including *btd* and *tll*) exclusively in T2NBs (Fig. S1B; Table S2). Enhancers in the NB lineage cohort are grouped into Pan-neural enhancers or T2 lineage (T2 lin) enhancers depending on chromatin accessibility and occupancy by Zld and Fru^C^ in T1NBs (Fig. 1C). Pan-neural enhancers show accessible chromatin and are bound by Zld and Fru^C^ and likely regulate gene expression in both NB lineages (Fig. S1B; Table S2). This category includes 1 enhancer from the *btd* locus active in T2NBs and 2 enhancers from the *D* and *opa* loci active only in INPs (Fig. S1C,D). Lack of Zld binding in T1NBs, which suggests inactivity, is used to define T2 lin enhancers. These include 6 enhancers from the *pnt, H6-like-homeobox* (*hmx*), *optix* (*opt*) and other loci that are active exclusively in T2NB lineages and 3 enhancers from the *erm* and *ham* loci active only in immature INPs (Fig. 1D,F; Fig. S1B; Table S2). We confirmed that T2 lin enhancers in the *erm* and *ham* loci indeed activate GFP reporter expression in immature INPs consistent with their endogenous protein expression patterns (Fig. 1E,G).^24,39,42^ T2 lin enhancers are prevalent in loci containing genes highly expressed in immature INPs suggesting that chromatin accessibility at cell-type-specific enhancers of genes essential for INP formation does not correlate with NB capability to generate INPs (Fig. 1A).

To correlate post-translational modifications of histones at cell-type-specific enhancers of genes essential for INP formation with the lack of INP progeny generated by T1NBs, T1NB-enriched chromatin was used to examine the patterns of active and repressive histone marks. Three histone modification profiles were established based on the combined pattern of active histone marks (H3K4me1, H3K4me3 and H3K27ac) and repressive histone mark (H3K27me3) ± 1kb from the center of the 21212 non-promoter regions with open chromatin in T2NBs. An active state (high H3K4me1/H3K27ac and low H3K27me3), a poised state (high H3K4me1/H3K27me3 and low H3K27ac) and an inactive state (low H3K4me1/H3K27ac/H3K27me3) (Fig. S1E). 10.1% of T2 lin enhancers and 14.5% of pan-neural enhancers display histone mark combination that is correlated with a poised state (Fig. 1H). The percentage of poised T2 lin enhancers and pan-neural enhancers that are associated with Ase^-^ and Ase^+^ immature INP genes (e.g. *Sp-1, erm, ham, Tfap2, D* and *Hmx*) was over-represented relative to randomized control in these two groups (Fig. 1H). Furthermore, the poised T2 lin and pan-neural enhancer category encompasses cell-type-specific enhancers that recapitulate the activation of endogenous *erm, ham, D* and *Hmx* expression (Fig. 1D-G). Thus, we conclude that post-translational modification of histones at enhancers of genes essential for INP formation correlates with the capability of NBs to generate INPs (Fig. 1A).

### Ase limits the capability to generate INPs in T1NBs

Accumulation of repressive histone mark H3K27me3 at cell-type-specific enhancers of genes essential for INP formation could contribute to their inactivity in T1NBs and limit INP generation. Yet, loss of Polycomb Repressive Complex 2 (PRC2) activity, the only enzymatic complex known to catalyze H3K27me3 deposition ^43,44^, had no apparent effect on GMC generation.^45^ Thus, it is unlikely that high levels of H3K27me3 alone maintain inactivity of these cell-type-specific enhancers. Poised enhancers can be activated by developmental TFs during lineage commitment in vertebrates.^46,47^ Btd is necessary for T2NBs to generate INPs.^20^ *btd* transcripts are highly enriched in T2NBs and immature INPs, and Btd protein binds T2 lineage enhancers of *erm* in T2NBs (Fig. 1D,F; Table S1). As such, lack of expression of T2 lineage-specific TFs including Btd could contribute to the inactivity of cell-type-specific enhancers that drive gene expression essential for INP formation in T1NBs.

High Btd expression can induce T1NBs to generate INPs, but with limited frequency.^20^ Thus, barriers limiting Btd-induced INP generation likely exist in T1NBs. We hypothesized that genes encoding these barrier proteins should be expressed exclusively in T1NBs or in T1NBs and INPs since these two cell types display similar transcriptomes and directly generate GMCs.^34–36^ Comparison of the transcriptome of wild-type T1NBs, T2NBs and INPs identified 37 genes that encode TFs and show transcripts enriched in T1NBs (Fig. 2A)^34–36^. We examined protein expression patterns for 35 of these genes and none were found to be uniquely expressed in all T1NBs at the protein level (Table S3). We next assessed 86 of 95 genes that encode TFs which show enrichment of transcripts in T1NBs and INPs but found *ase* and *pros* were the only genes whose proteins are detected in T1NBs but not in T2NBs (Table S3). *pros* was excluded from further analyses because Pros functions in GMCs to promote cell cycle exit and differentiation.^15–17,48^ We conclude that Ase is a candidate regulator that limits INP generation in T1NBs.

**Figure 2.**
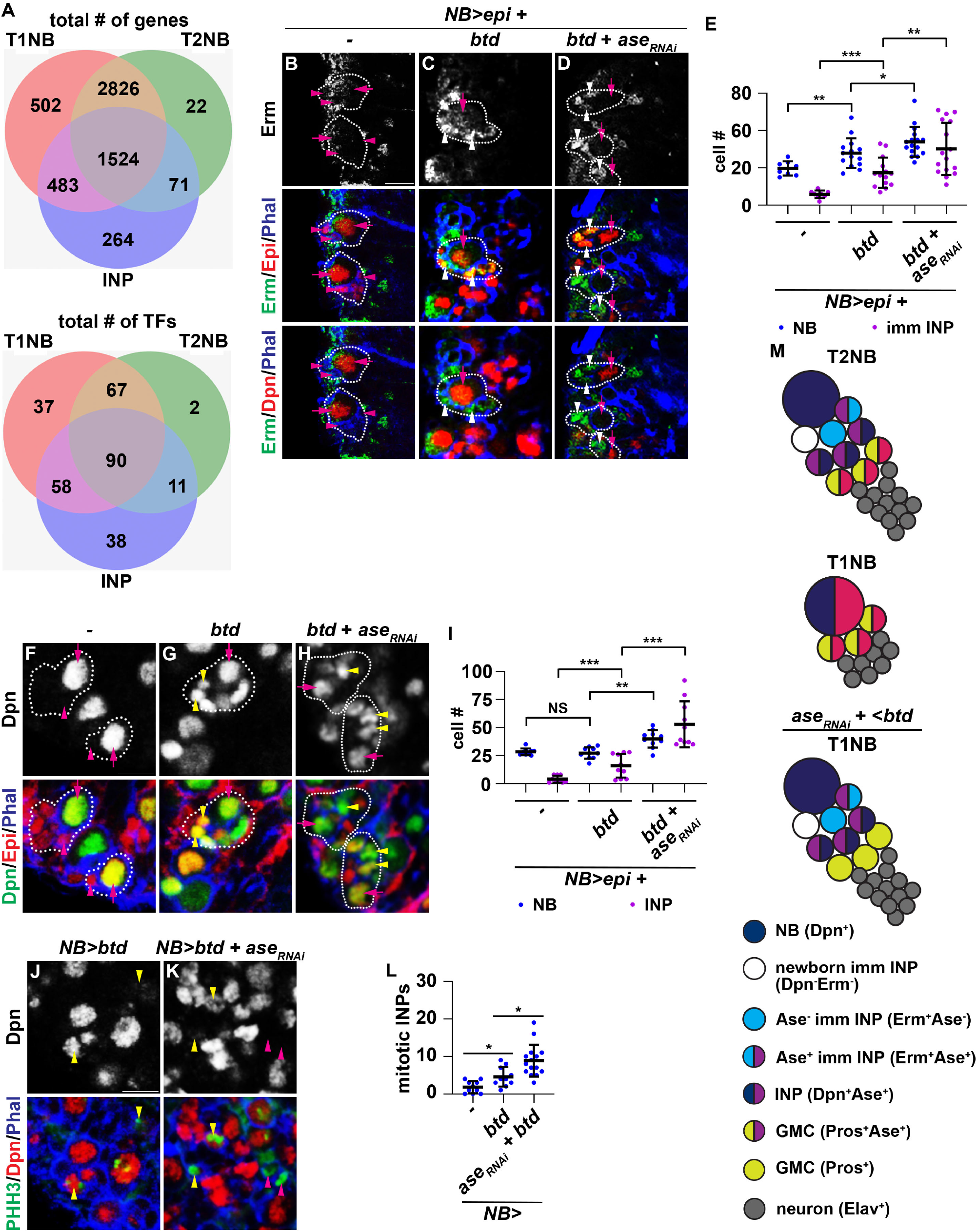
Ase limits the capability of T1NBs to generate INPs induced by Btd. (A) Single-cell transcriptomic comparisons of enriched transcripts (all transcripts – top; TFs only -bottom) across T1NBs, T2NBs and INPs from published single-cell mRNA sequencing datasets of wild-type NB lineages.^34–36^ (B-D) Images showing that *ase* knockdown with pan-NB driver (*Wor-Gal4*) increases the efficiency of Btd mis-expressing T1NBs, located on the ventral surface of the larval brain, to generate immature INPs (small epitope^+^Erm^+^Dpn^-^ cells). (E) Quantification of number of ventral NBs and ventral immature INPs showing knockdown of *ase* increased number of immature INPs compared to NBs upon Btd mis-expression. (F-H) Images showing that *ase* knockdown with pan-NB driver increases the efficiency of Btd mis-expressing T1NBs to generate INPs (small epitope^+^Dpn^+^ cells). (I) Quantification of number of ventral NBs and ventral INPs showing knockdown of *ase* increased number of INPs compared to NBs upon Btd mis-expression. (J-K) Images showing that *ase* knockdown with pan-NB driver increases the efficiency of Btd mis-expressing T1NBs to generate mitotic INPs (small PHH3^+^Dpn^+^ cells). (L) Quantification of number of mitotic INPs found on the ventral side of the brain showing knockdown of *ase* increased number of mitotic INPs compared to Btd mis-expression. (M) Diagram summarizing the expression patterns of cell identity markers in T1NBs, T2NBs, and Btd mis-expressing T1NBs combined with *ase* knockown and their progeny. T1NBs lacking Ase and mis-expressing Btd generate Erm^+^ immature INPs that mature into proliferative INPs. White dotted line marks NB lineage. Magenta arrow: T1NB. Magenta arrowhead: GMC. White arrowhead: Erm^+^ immature INP. Yellow arrowhead: INP. Scale bar: 10 µm. P-value; NS: Not significant,, *<.05., **<.005., ***<.0005. ****<.00005.

If Ase functions as a barrier to INP generation, reducing *ase* function should allow Btd to more efficiently induce T1NBs to generate INPs. We used a *NB-Gal4* driver (*Wor-Gal4*) to induce *UAS-btd::myc* expression in all NBs and assessed the identity of their newly generated progeny marked by the perdurance of the Myc epitope expression. We focused on the ventral surface of larval brains which contains exclusively T1NBs under wild-type conditions.^15–17^ Control brains that expressed a Myc epitope reporter transgene contain NBs that produce GMCs that do not express the immature INP marker Erm (Fig. 2B,E). In contrast, high levels of Btd expression in T1NBs led to generation of small Dpn^-^Erm^+^ cells (induced immature INPs) which are found in direct contact with the NB (Fig. 2C, E). Importantly, after reducing *ase* function in tandem with high levels of Btd, the number of induced immature INPs produced significantly increased (Fig. 2D, E). To confirm differentiation of these induced immature INPs into INPs we used the following 2 criteria: reactivation of Dpn expression to validate INP identity, and positive phospho-histone H3 (PHH3) to confirm mitotic potential. None of the small Myc^+^ cells in control brains express Dpn (Fig. 2F, I). Btd mis-expressing brains contain 16±11 small Myc^+^ cells that express Dpn, while knocking down *ase* function increased the total number of small Myc^+^Dpn^+^ cells to 53±21 per brain lobe (Fig. 2G-I). We detected 5±3 small Myc^+^,Dpn^+^ cells that are mitotically active marked by PHH3 in brains that expressed elevated levels of Btd (Fig. 2J,L). Reducing *ase* function significantly increased the total number of small Myc^+^Dpn^+^PHH3^+^ cells per lobe (Fig. 2K,L). These results support a role for Ase in limiting the ability of T1NBs to generate INPs (Fig. 2M). Thus, we conclude that Ase functions as a barrier to INP generation in T1NBs.

### Elevated Ase expression promotes T2NBs to generate GMCs via bypassing INPs

We tested whether Ase is sufficient to limit the ability of T2NBs to generate INPs by elevating Ase expression. Instead of relying on the lack of Ase expression for identifying T2NBs in this experiment, we overexpressed a *UAS-GFP* transgene driven of a T2NB-specific Gal4 driver (*tll-Gal4*) to positively mark T2NB lineages.^23,24^ 83% of T2NBs in wild-type brains produce Erm^+^ immature INPs which are found in direct contact with the NB (Fig. 3A,C). In contrast, elevating Ase expression drastically reduced the percentage of T2NBs that produce immature INPs with most NBs producing Erm^-^ cells. This result suggests that Ase inhibits T2NBs from generating immature INPs (Fig. 3B,C). Consistent with a reduced number of immature INPs, high levels of Ase expression led to significantly fewer INPs marked by Dpn expression per lobe (20±8 per lobe) than wild-type brains (129±15 per lobe) (Fig. S2A,B,D). Similar to Ase, mis-expression the human homolog Ascl1 led to significantly fewer INPs per brain lobe (24±7 per lobe) and T2NBs in these brains are directly surrounded by small Dpn^-^Ase^+^ cells that are likely GMCs (Fig. S2C,D). We could unambiguously quantify INPs in this experiment because the antibody we used against fly Ase does not cross-react with human Ascl1. We further confirmed the identity of these small Dpn^-^Ase^+^ cells using nuclear localization of Pros, an indicator of GMC identity. We found that 98% of T2NBs upon Ase mis-expression were found with GMCs in direct contact with the NB whereas wild-type T2NBs had GMCs distal to NB (Fig. 3D-F). Altogether, these results suggest that T2NBs expressing elevated levels of Ase generate GMCs instead of INPs.

**Figure 3.**
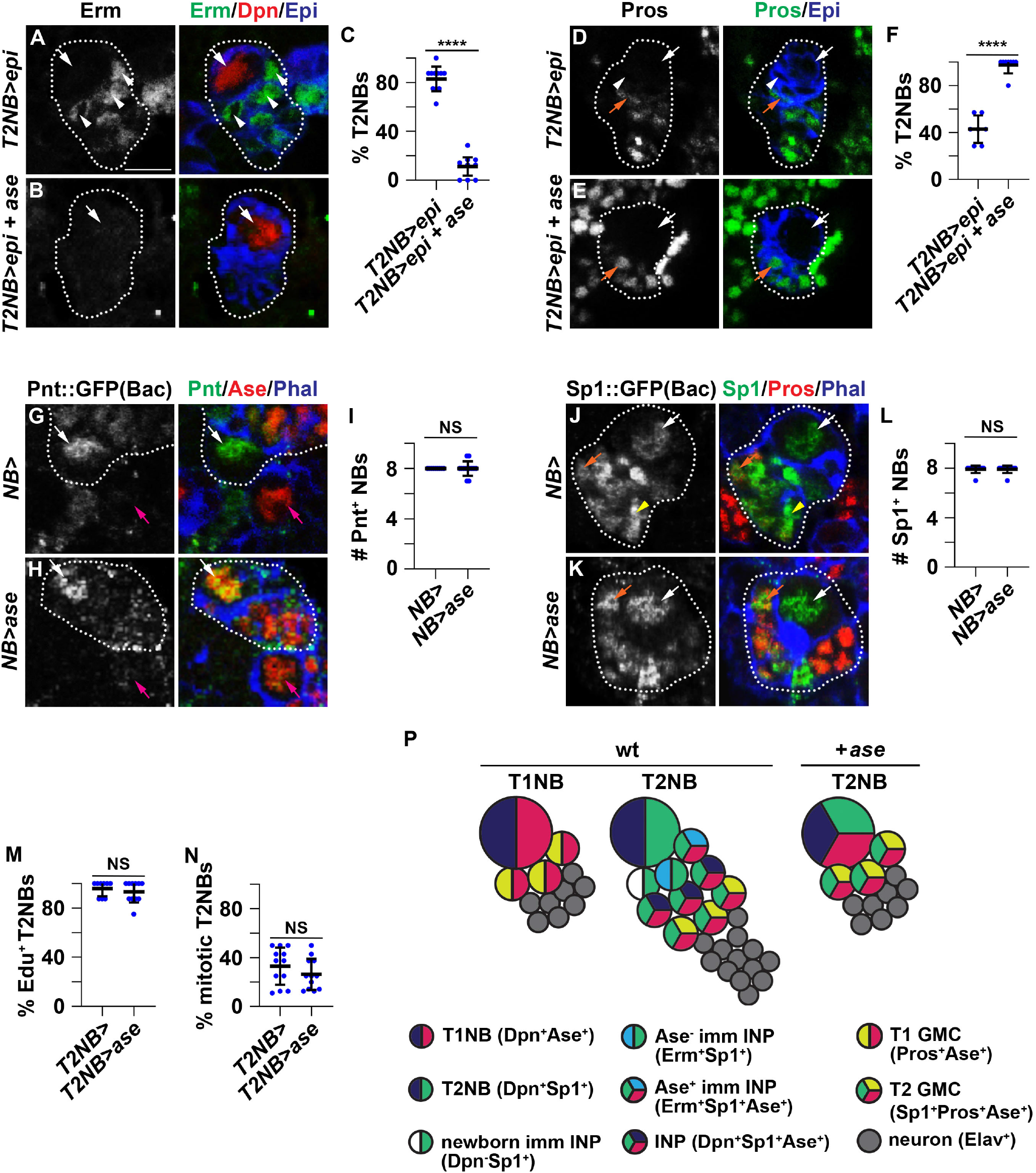
Elevated Ase expression promotes T2NBs to generate GMCs via bypassing INPs. (A-B) Images showing that mis-expressing Ase with a T2NB driver (*Tll-Gal4*) results in T2NBs no longer generating immature INPs. A wild-type T2NB marked by GFP expression is always in contact with immature INP progeny that express Erm. An Ase mis-expressing T2NB is in contact with progeny lacking Erm expression. (C) The percentage of Ase mis-expressing T2NBs surrounded by Erm-positive progeny is significantly lower than control T2NBs. (D-E) Images showing that mis-expressing Ase with a T2NB driver results in T2NBs generating GMCs directly. GMCs identified by Pros expression are usually located at least one-cell away from a control T2NB but are found in contact with the Ase mis-expressing T2NBs. (F) The percentage of Ase mis-expressing T2NBs in direct contact with GMCs is significantly higher than control T2NBs. (G-H) Images showing T2NBs mis-expressing Ase using a pan-NB driver (*Wor-Gal4*) still maintain T2NB-specific marker Pnt expression. (I) The number of NBs demonstrating high Pnt expression is not significantly different between control versus Ase mis-expression. (J-K) Images showing T2NBs mis-expressing Ase using a pan-NB driver still maintain T2NB-specific marker Sp1 expression. (L) The number of NBs demonstrating Sp1 expression is not significantly different between control versus Ase mis-expression. (M-N) The percentage of control versus Ase mis-expressing T2NBs displaying positive EdU incorporation or PHH3 is statistically indistinguishable. (P) Diagram demonstrating how T2NBs mis-expressing Ase bypass INPs to generate GMCs while maintaining their lineage identity. White dotted line marks NB lineage. White arrow: T2NB. White arrowhead: Erm^+^ immature INP. Yellow arrowhead: INP. Magenta arrow: T1NB. Orange arrow: GMC. Scale bar: 10 µm. P-value; NS: Not significant, ****<.00005.

To test if Ase induced T2NBs to generate GMCs by converting their identity to a T1NB state, we used four methods to assess T2NB identity in control brains carrying a NB-specific Gal4 driver alone and in brains that expressed elevated levels of Ase (Fig. S2E). The *pnt::GFP(Bac)* transgene recapitulates endogenous PntP1 expression in larval brains including T2NB-specific expression.^23^ The number of Pnt::GFP^+^ NBs appears statistically indistinguishable between control and elevated Ase samples (Fig. 3G-I). Due to lack of reliable antibodies or reporter transgenes, we performed single-molecule fluorescent *in situ* hybridization (sm-FISH) to assess *btd* transcript patterns. T2NBs were marked by RFP driven by a T2NB-specific Gal4 and *btd* transcript levels were normalized against transcripts of the ribosomal gene *mRpl15*. The relative levels of *btd* transcripts in T2NBs are not significantly different between control and elevated Ase samples (Fig. S2F-H). The *tll::GFP(Bac)* transgene mimics endogenous Tll expression in larval brains including higher levels in T2NBs compared to T1NBs.^23,24^ The number of NBs expressing high levels of Tll per lobe appears statistically indistinguishable between control and elevated Ase samples (Fig. S2I-K). The *Sp1::GFP(Bac)* transgene is expressed throughout the entire T2 lineage.^24,34^ The number of Sp1::GFP^+^ NBs per lobe appears statistically indistinguishable between control and elevated Ase samples (Fig. 3J-L). GMCs in direct contact with T2NBs that expressed high levels of Ase maintained Sp1::GFP expression, indistinguishable from GMCs generated by T2NBs in control brains (Fig. 3J,K). Together, these results support a model that high levels of Ase induce T2NBs to generate GMCs without altering their lineage identity.

To exclude the possibility that changes in cell proliferation led T2NBs to generate GMCs instead of INPs, we examined Edu incorporation and the mitotic marker PHH3 in T2NBs that expressed high levels of Ase compared to wild-type samples. We observed no changes in Edu incorporation upon Ase mis-expression, with all T2NBs taking up Edu after a three-hour feeding similar to control samples (Fig. 3M). Furthermore, the percentage of PHH3^+^ T2NBs in brains that expressed elevated levels of Ase in T2NBs is indistinguishable from control (Fig. 3N). Thus, it is unlikely that high levels of Ase induces T2NBs to generate GMCs instead of INPs by altering cell proliferation. We conclude that high levels of Ase promote T2NBs to generate GMCs via bypassing INPs (Fig. 3P).

### Ase functions through Prospero to promote generation of GMCs by T2NBs

A key first step to identify the downstream-effectors that mediate Ase-induced reprogramming of T2NBs to generate GMCs is to define whether Ase limits INP generation by activating or repressing gene transcription. We took advantage of a previously published *UAS-ase::VP16* transgene ^32^ that encodes a constitutive transcriptional activator form of Ase and mis-expressed Ase::VP16 in T2NBs. The Ase antibody does not recognize Ase::VP16, nor does Ase::VP16 appear to promote expression of endogenous Ase, allowing us to unambiguously identify T2NBs and INPs (Dpn^+^Ase^+^). Ase::VP16 mis-expression in T2NBs phenocopies the effect of full-length Ase and Ascl1 mis-expression showing a significant reduction in the number of INPs per lobe without affecting the expression of T2NB-specific markers including Sp1 (Fig. S3A-F). Importantly, T2NBs mis-expressing Ase::VP16 are also surrounded by GMCs (Dpn^-^Ase^+^) instead of INPs (Fig. S3B). These results indicate that Ase induces T2NBs to generate GMCs by activating gene expression.

To identify the downstream-effectors of Ase in inducing GMC generation, we used a previously established genetic strategy to enrich for T1NBs (Fig. S1A).^20,49^ We confirmed that the transcriptome of enriched T1NBs is highly similar to wild-type T1NBs determined by sc-RNA sequencing (Fig. S1A).^35^ T1NB-enriched chromatin was used to identify 2993 active neurogenic enhancers defined as non-promoter *cis*-regulatory regions that display accessible chromatin and are bound by Zld and Fru^C^ (Fig. 4A). 2589 of these enhancers are bound by Ase in T1NBs. Because Ase is highly expressed in T1NBs and INPs, gene transcripts that are nearest to Ase-bound neurogenic enhancers and are enriched in T1NBs alone or in T1NBs and INPs are considered as candidate Ase downstream-effectors (Table S3). 18 of these 113 genes encode TFs, with *ase* and *pros* being the only two genes whose protein products are specifically expressed in T1NBs and INPs (Table S4). We focused on *pros* and confirmed that high levels of Ase were sufficient to induce *pros* transcript enrichment in T2NBs (Fig. 4B-D). Thus, *pros* is an excellent candidate downstream-effector of Ase.

**Figure 4.**
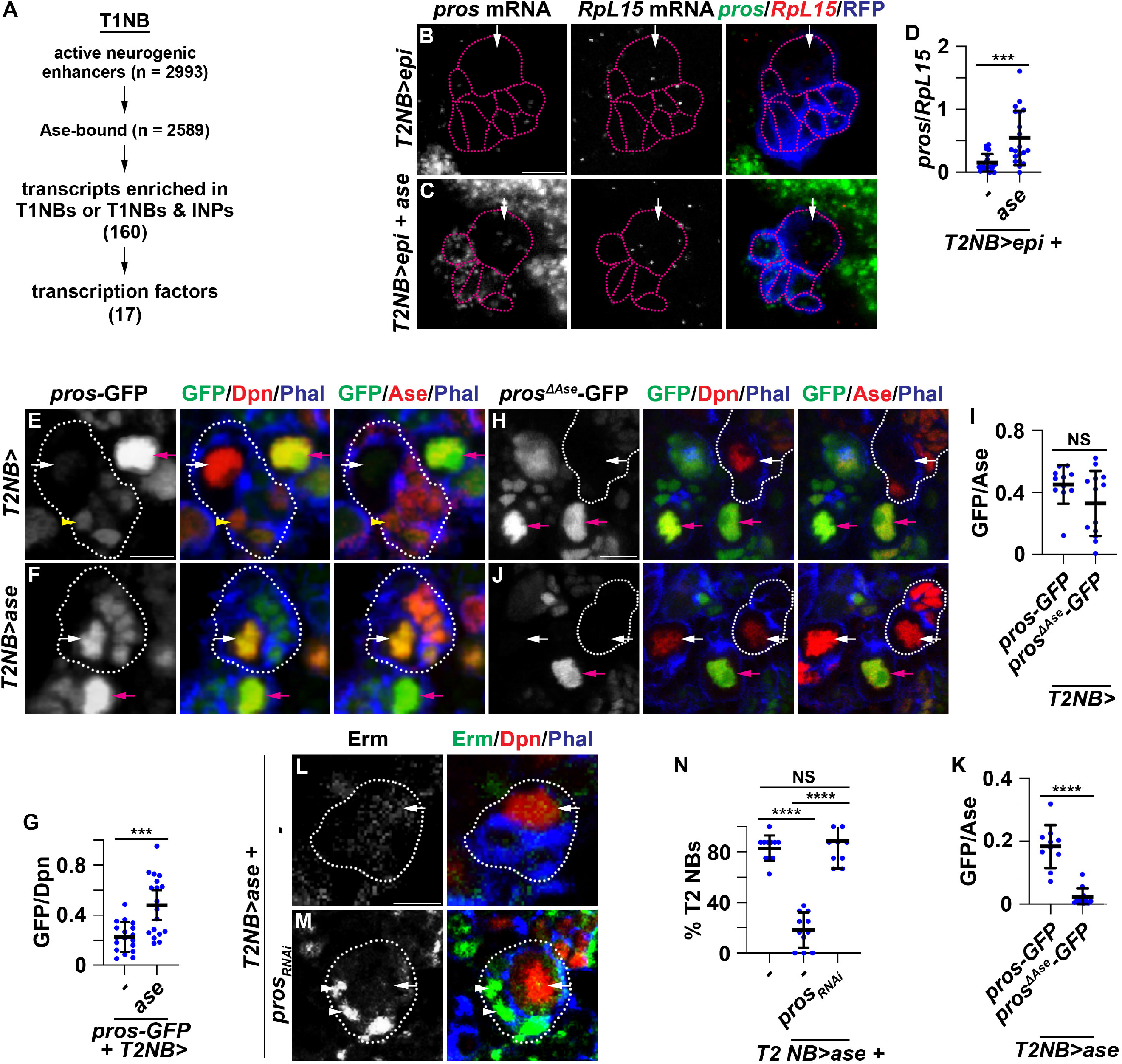
Ase functions through Pros to promote generation of GMCs by T2NBs. (A) A flow chart for identifying candidate Ase downstream-effector genes that encode transcription factors. (B,C) Images showing Ase mis-expression is sufficient to induce *pros* transcription in T2NBs. *pros* transcripts are undetectable in control T2NBs but robustly detected in Ase mis-expressing T2NBs. *RpL15* transcripts serve as an internal control. (D) Levels of *pros* transcripts relative to *RpL15* transcripts in Ase mis-expressing T2NBs are significantly higher than control T2NBs. (E,F) The expression of a GFP reporter controlled by enhancer 3 from the *pros* locus bound by Ase can be activated in T2NBs by Ase mis-expression. *pros*^*enh3*^-GFP (abbreviated as *pros*-GFP) is expressed in T1NBs and INPs but not in T2NBs in wild-type brains mimicking endogenous Ase expression. *pros*-GFP expression is detected in T2NBs mis-expressing Ase. (G) Levels of *pros*-GFP expression relative to Dpn expression are significantly higher in T2NBs mis-expressing Ase than control T2NBs. (H) Removing all Ase-binding sites in the enhancer 3 from the *pros* locus has no effect on its cell-type-specific expression pattern. The *pros*^*enh3ΔAse*^-GFP (abbreviated as *pros*^*ΔAse*^-GFP) reporter transgene lacks all Ase-binding sites and is expressed in T1NBs and INPs but not in T2NBs. (I) Levels of *pros*^*ΔAse*^-GFP expression relative to background Ase staining are statistically indistinguishable from *pros*-GFP expression. (J) The *pros*^*ΔAse*^-GFP reporter transgene is not activated in T2NBs upon Ase mis-expression. (K) Levels of *pros*^*ΔAse*^-GFP relative to Ase staining are significantly lower than *pros*-GFP expression. (L-M) Images showing that knocking down *pros* function in Ase mis-expressing T2NBs restores their ability to generate immature INPs. An Ase mis-expressing T2NB is surrounded by progeny lacking Erm expression but become surrounded by immature INPs marked by Erm expression when *pros* function is knocked down. (N) The percentage of Ase mis-expressing T2NBs surrounded by Erm^+^ immature INPs is restored to a level indistinguishable from wild-type T2NBs following *pros* knockdown. White dotted line marks NB lineage. White arrow: T2NB. White arrowhead: Erm^+^ immature INP. Yellow arrowhead: INP. Magenta arrow: T1NB. Scale bar: 10 µm. P-value; NS: Not significant, ****<.00005, ***<.0005.

We assessed enhancer activity under the control of each of three putative Ase-bound enhancers in the *pros* locus (Fig. S3G). GFP expression driven by enhancer 1 or 2 was detected in T2NBs in addition to T1NBs and INPs or detected only in a small subset of T1NBs suggesting that these two regulatory elements are not activated by Ase (Fig. S3G,H). In contrast, the expression of *pros-GFP* reporter driven by enhancer 3 was detected in T1NBs and INPs but not T2NBs, a pattern consistent with endogenous Ase expression patterns (Fig. 4E, S3G). Elevated levels of Ase were also sufficient to activate *pros-GFP* expression in T2NBs (Fig. 4F,G). Enhancer 3 contains 4 putative Ase-binding sequences.^50^ The expression pattern and intensity of the *pros*^*ΔAse*^*-GFP* reporter, which carries deletion of all 4 Ase-binding sites, appeared indistinguishable from *pros-GFP* in control brains (Fig. 4H,I). Most importantly, elevated levels of Ase cannot activate *pros*^*ΔAse*^*-GFP* expression in T2NBs (Fig. 4J,K). Thus, we conclude Ase functions through enhancer 3 to regulate *pros* expression.

If Pros mediates Ase-induced reprogramming of T2NBs to generate GMCs, knocking down *pros* function upon Ase misexpression should restore INP generation. Indeed, reducing *pros* function in T2NBs that mis-express Ase restored the generation of immature INPs marked by Erm expression (Fig. 4L-N). These immature INPs were able to differentiate into INPs marked by *odd paired* (*opa*) transcripts and Dpn protein (S3I-L). Thus, we conclude that Ase functions through *pros* to reprogram T2NBs to generate GMCs.

### Prospero reprograms immature INPs to bypass an INP state and adopt a GMC fate

Although *pros* is required for Ase-induced reprogramming of T2NBs to generate GMCs (Fig. 4L,M, S3I-K), nuclear Pros protein first becomes detectable in immature INPs in direct contact with these NBs rather than in the NB (Fig. 3E, S2C). This result suggests that high levels of Ase expression in T2NBs leads to activation of the GMC genetic program in immature INPs. This hypothesis predicts that GMCs are more closely related to immature INPs than NBs or INPs. We used published single-cell mRNA sequencing datasets of wild-type larval brains to find the transcriptomic similarity between NBs, immature INPs, INPs, GMCs and neurons, using the Euclidean distance between average gene expression of each cell-type (Fig. 5A).^34–36^ GMCs in both NB lineages are the most closely related cell type to each other, while Ase^-^ and Ase^+^ immature INPs are most closely related to one another (Fig. 5A). Consistent with our hypothesis, the next most closely related cell type to GMCs is Ase^-^ immature INPs followed by Ase^+^ immature INPs (Fig. 5A). We tested whether elevated Pros levels can activate the GMC genetic program in immature INPs by expressing a *UAS-pros* transgene under the control of an immature INP Gal4 driver (*erm*-GAL4) (Fig. 5B). Brains expressing elevated levels of Pros contained significantly fewer INPs per lobe than control brains (Fig. 5C-E). While T2NBs in control brains were surrounded primarily by INPs (Dpn^+^Ase^+^), cells in direct contact with T2NBs in brains that expressed elevated levels of Pros appeared to be GMCs (Dpn^-^ Ase^+^) (Fig. 5C,D). These data suggest that Pros expression can activate the GMC genetic program in immature INPs.

**Figure 5.**
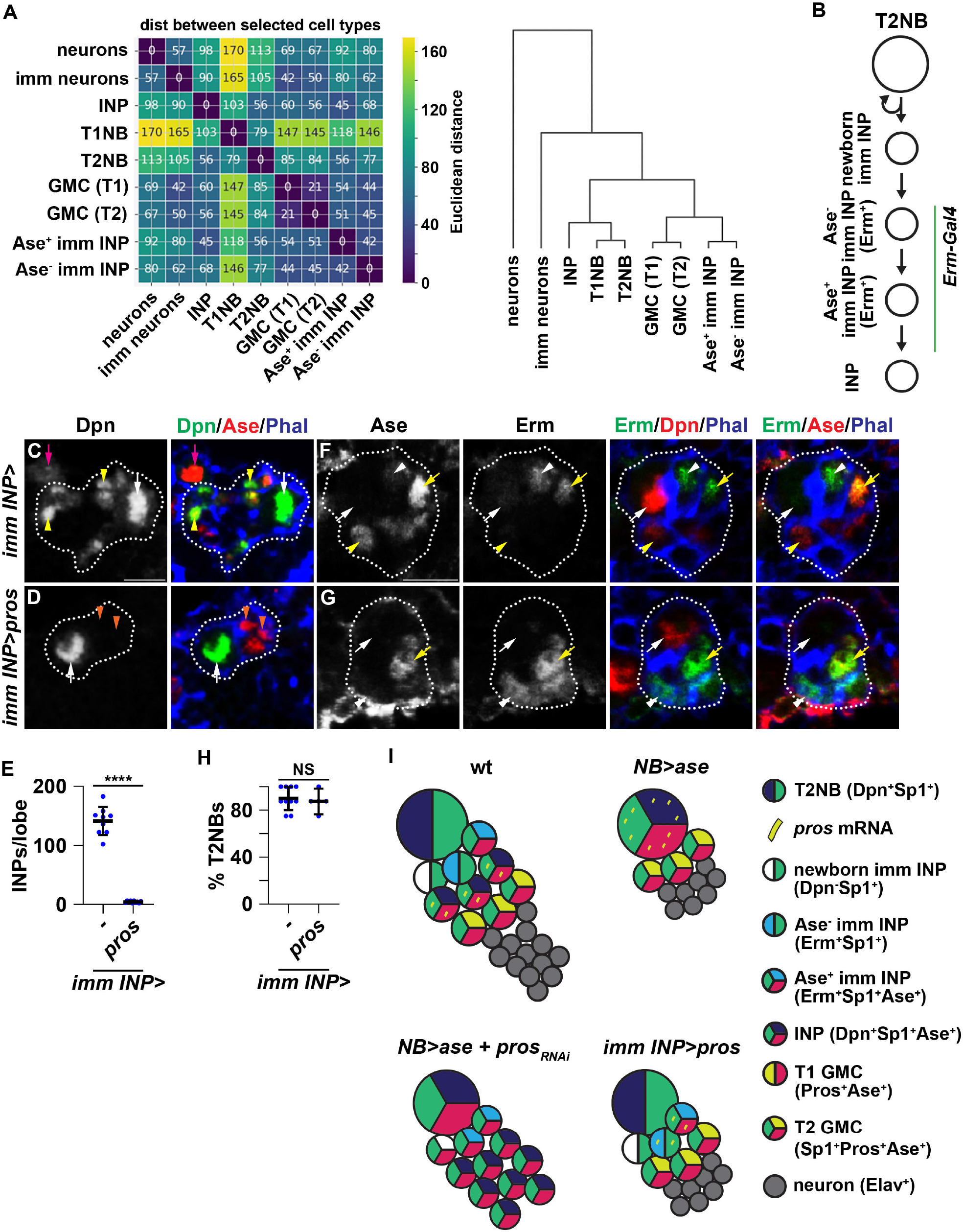
Pros reprograms immature INPs to bypass an INP state and adopt a GMC fate. (A) The Euclidean distance between average gene expression of NBs, immature INPs, INPs, GMCs and neurons using published single-cell mRNA sequencing datasets of wild-type larval brains.^34–36^ GMCs from the T1 and T2 lineages are highly similar to immature INPs. Dendrogram based on Pearson correlation summarizes the degree of similarity between sell types. (B) Diagram depicting at what stage of immature INP *Erm-Gal4* is active in the T2 lineage. (C-D) Images showing Pros mis-expression in immature INPs by *Erm-Gal4* promotes premature adoption of a GMC fate. A wild-type T2NB lineage contains INPs (small Dpn^+^Ase^+^ cells) whereas a T2NB lineage that mis-expresses *pros* in immature INPs only contains small Dpn^-^Ase^+^ progeny (E) Quantification showing the number of INPs per brain lobe where Pros is mis-expressed in immature INPs is drastically lower than wild-type brain lobe. (F-G) Images showing Pros mis-expression in immature INPs does not prevent specification of Erm^+^ immature INPs or Ase^+^ immature INPs. (H) Quantification showing the fraction of T2NBs producing Erm^+^ immature INPs upon Pros mis-expression in immature INPs not significantly different from wild-type T2NBs. (I) A schematic summarizing the data that suggest Pros mediates Ase-induced bypassing of INP production to generate GMCs by T2NBs. White dotted line marks NB lineage. White arrow: T2NB. White arrowhead: Erm^+^/Ase^-^ immature INP. Yellow arrow: Ase^+^ immature INP. Yellow arrowhead: INP. Magenta arrow: T1NB. Orange arrow: T2 lineage GMC. Scale bar: 10 µm. P-value; NS: Not significant, ****<.00005

We tested whether Pros activates the GMC genetic program in immature INPs in part by overriding immature INP gene expression. Newborn immature INPs transition to Ase^-^ immature by upregulating Erm expression 90 minutes following asymmetric division of the T2NB.^27,40^ As such, Erm protein expression is the earliest marker that indicates the onset of differentiation in a newborn immature INP. Pros mis-expression in immature INPs had no effect on endogenously expressed Erm protein and upregulation of Ase expression (Fig. 5F-H). Thus, we conclude that Pros-induced reprogramming of immature INPs into GMCs occurs through superimposition of the GMC genetic program upon endogenously expressed immature INP genes (Fig. 5I).

### Reprogramming tumor NB progeny fate bypasses differentiation blockades and halts tumor growth

The ability to control the generation of specific TAP subtypes could provide a strategy to bypass differentiation blockades in TAPs that leads to tumorigenesis. The RNA-binding protein Brain Tumor (Brat) promotes the onset of differentiation in newborn immature INPs, and loss of *brat* function blocks immature INP differentiation to INPs, resulting in a transformation into premalignant T2NBs.^15–17,51–55^ A subset of these premalignant T2NBs bypass cell cycle exit and immortalize into tumor NBs in adult *brat*-mutant brains.^56,57^ Adult *brat*-mutant tumor NBs lack Ase expression suggesting that they maintain T2NB identity and likely generate immature INPs that are stalled in differentiation and revert to tumor NBs (Fig. 6A; Fig. S4A).

**Figure 6.**
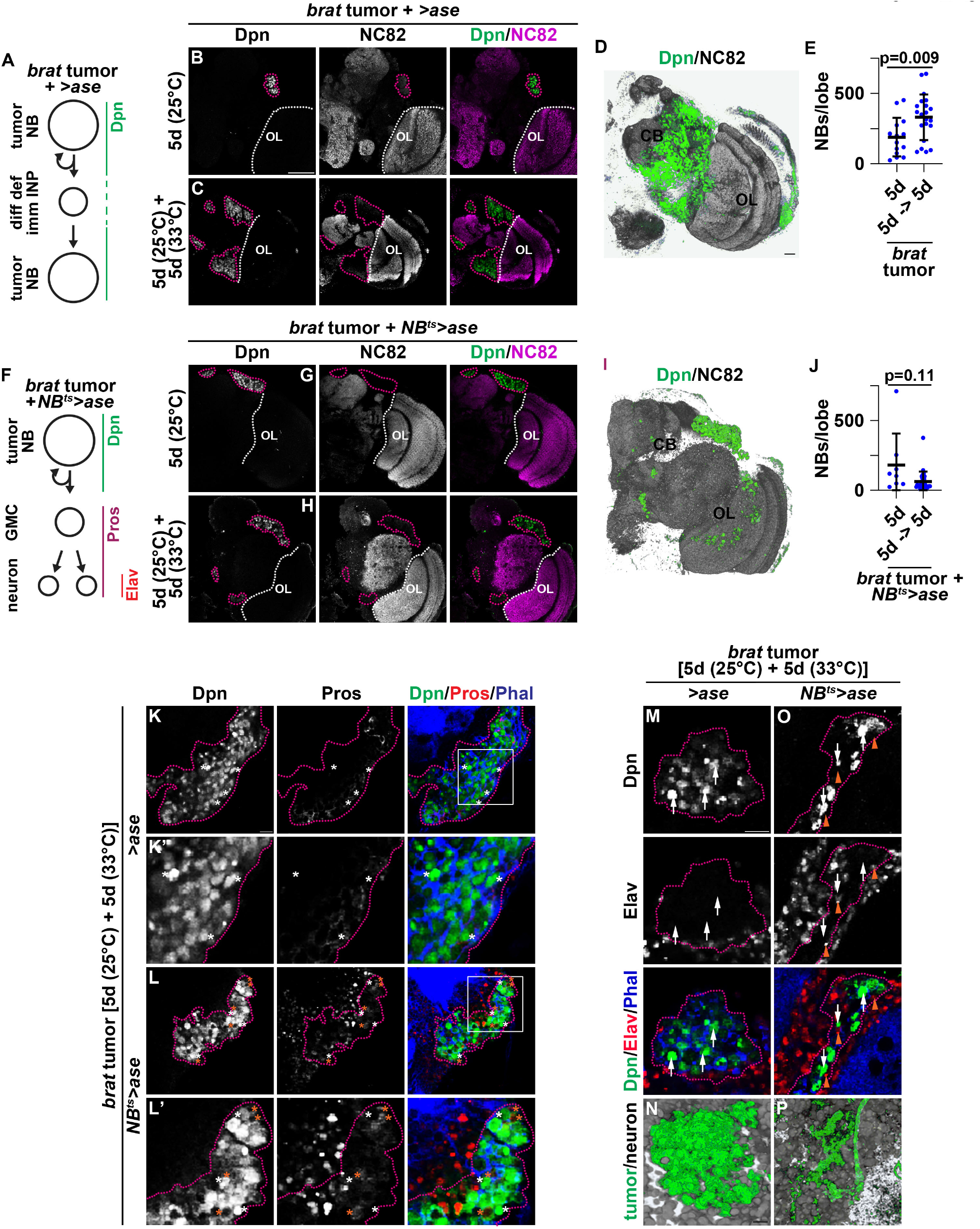
Reprogramming tumor NBs to generate GMCs halts tumor growth driven by defective INPs. (A) Diagram showing that upon loss of *brat* function, immature INPs fail to differentiate into INPs and revert back to tumor NBs. (B-C) Images showing tumor NBs continually grow in adult fly brains in a *brat*-, *patr-1* hypomorphic background. Tumor cells are marked by Dpn expression, and neurons are marked by NC82 expression. Scale bar: 50 µm. (D) 3-D reconstruction of an adult brain showing tumor mass (pseudo colored in green) 10 days after eclosion. Scale bar: 10 µm. (E) Quantification showing that the number of tumor NBs (Dpn^+^ cells) increases over time. (F) Diagram showing that mis-expression of Ase upon loss of Brat function will produce progeny that bypass immature INP state and restore generation of differentiated progeny marked with Pros (GMCs) or Elav (neurons). (G-H) Images showing tumor NBs stop growing or may become ablated in adult fly brains upon Ase mis-expression in a *brat*-, *patr-1* hypomorphic background. (I) 3-D reconstruction of an adult brain showing tumor mass (pseudo colored in green) 10 days after eclosion is much smaller in the Ase mis-expressing samples. (J) Quantification showing that the number of tumor NBs (Dpn^+^ cells) upon Ase mis-expression is not significantly different and is likely decreasing over time. (K-L) Images showing tumor NBs are induced to produce GMC progeny (Pros^+^) upon Ase mis-expression in adult fly brains in a *brat*-, *patr-1* hypomorphic background. Scale bar: 20 µm. (M,O) Images showing tumor NBs are induced to produce neurons progeny (Elav^+^) upon Ase mis-expression in adult fly brains in a *brat*-, *patr-1* hypomorphic background. White dotted line marks OL boundary. Magenta dotted line marks tumor boundary. Scale bar: 20 µm. (N,P) 3-D reconstruction of an adult brain showing tumor mass (pseudo colored in green) is invaded by neurons (pseudo colored gray) upon Ase misexpression. Scale bar: 20 µm.

If tumor cells-of-origin in adult *brat*-mutant brains are indeed immature INPs stalled in differentiation, Ase-induced reprogramming of immature INPs into GMCs should limit tumor expansion. Brat promotes degradation of gene transcripts that encode proteins inhibiting the onset of differentiation in immature INPs.^37,58^ Protein associated with topo II related-1 (Patr-1) is a component of the RNA decapping protein complex, which contributes to mRNA decay.^59^ Brat and Patr-1 function together to promote immature INP differentiation to INPs.^55^ Larval brains carrying *brat* and *patr-1* hypomorphic alleles (*brat*^*hypo*^,*patr-1*^*hypo*^) display a moderate premalignant T2NB phenotype originating from immature INPs stalled in differentiation (Fig. S4B-D,F,G). High levels of Ase expression significantly suppressed the premalignant T2NB phenotype in *brat*^*hypo*^,*patr-1*^*hypo*^ larval brains (Fig. S4D-F). Wild-type adult brains do not contain NBs, whereas *brat*^*hypo*^,*patr-1*^*hypo*^ adult brains displayed detectable tumor NBs marked by Dpn expression with high penetrance at eclosion (Fig. S4G). We used *brat*^*hypo*^,*patr-1*^*hypo*^ animals that carry a *UAS-ase* transgene alone as control animals that receive no treatment. *brat*^*hypo*^,*patr-1*^*hypo*^ animals that carried a heat-inducible *NB-Gal4* driver and a *UAS-ase* transgene were experimental animals where Ase expression can be induced in tumor NBs shifting from 25°C to 33°C. Tumor masses were identified using Dpn to mark NBs and NC82 to mark the neuropil. Tumor masses were clearly visible in controls 5 days post-eclosion at 25°C and increased their volume and total number of tumor NBs following additional 5 days of incubation at 33°C (Fig. 6B-E, Supplementary Movie 1). Although tumor masses were also visible in experimental animals 5 days post-eclosion at 25°C, tumor NB number and tumor burden failed to increase upon induction of Ase expression for 5 days (Fig. 6F-J, Supplementary Movie 2). These results demonstrate that Ase-induced reprogramming of immature INPs to adopt a GMC fate can bypass differentiation blockades in immature INPs and halt tumor growth.

We further examined whether Ase expression indeed halts *brat*^*hypo*^,*patr-1*^*hypo*^ tumor expansion by reprogramming tumor NBs to generate GMCs marked by nuclear Pros. We also assessed if these GMCs were able to differentiate into neurons by utilizing the neuron marker Elav. The *brat*^*hypo*^,*patr-1*^*hypo*^ control brains consists of mostly tumor NBs marked by Dpn expression and very few GMCs and neurons (Fig. 6K,M,N). We did not detect any GMCs and neurons immediately adjacent to tumor NBs (Fig. 6K’,M). In contrast, in *brat*^*hypo*^,*patr-1*^*hypo*^ brains that had Ase expression induced, it could be observed that GMCs and neurons were interspersed among tumor NBs (Fig. 6L,O,P). Importantly, we could easily detect GMCs and neurons located adjacent to tumor NBs (Fig. 6L’,M). This result is consistent with GMCs surrounding T2NBs that expressed high levels of Ase in larval brains (Fig. 3E). Thus, we conclude that the Ase-instilled barrier to generate INPs can halt tumor expansion by reprogramming tumor NBs to generate GMCs via bypassing stalled INPs that are the cells-of-origin in *brat*-mutant brains.

## Discussion

Despite significant advance in our understanding of the genomic landscape and transcriptome of various cell types in the developing vertebrate cortex ^9,60^, how radial glia decide to generate specific TAP subtypes remain unknown. We investigated how NBs decide to generate INPs instead of GMCs during fly larval brain neurogenesis. The capability to generate INPs was defined by chromatin accessibility at cell-type-specific enhancers of genes that drive INP formation in the T2 lineage (Fig. 1C). Surprisingly, these enhancers are maintained in a poised state in T1NBs despite never generating INPs in their lifetime (Fig. 7A). Thus, both NB types are competent to generate INPs. The mutually exclusive expression and functions of Btd and Ase/Pros maximizes the efficiency of INP generation in T2NBs and GMC production in T1NBs in wild-type brains (Fig. 7B). Because cell-type-specific enhancers of genes essential for INP formation are maintained in a poised state in T1NBs, endogenously expressed Ase safeguards GMC generation by constraining INP production induced by aberrant expression of T2NB TFs such as Btd (Fig. 7B). Elevating Ase expression in T2NBs functions through Pros to reprogram immature INPs into GMCs bypassing the INP state. Our study suggests that a double-assurance mechanism in which dynamic superimposition of lineage-specific activators (Btd) upon basal-state determinants (Ase/Pros) defines NB capability to generate INPs or GMCs.

**Figure 7.**
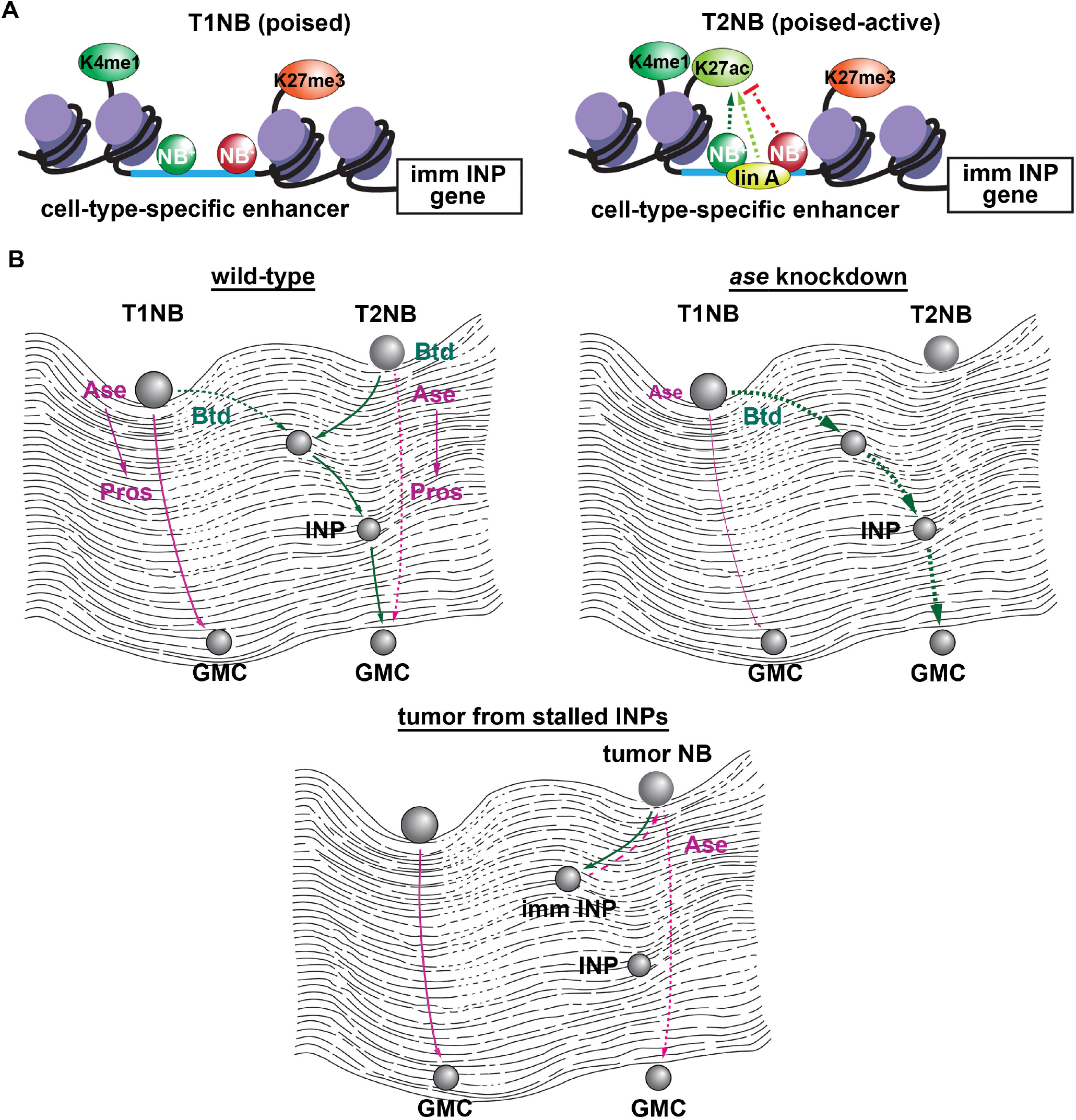
The capability to generate INPs is inherent to both NB types but activated exclusively in T2NBs. (A) Enhancers associated with genes that promote INP generation are maintained in a poised state in T1NBs but are in a poised-activate state in T2NB through binding of lineage specific TFs. NB^+^: NB-specific transcriptional activator, NB^-^: NB-specific transcriptional repressor, Lin A: Lineage-specific transcriptional activator. (B) Chromatin landscape diagram showing that both T1 and T2NBs are capable to generate INPs or GMCs directly based on cues from TFs. Stoichiometric superimposition of basal-state transcription factors (Ase and Pros) and lineage-specific transcription factor (Btd) defines NB decision to generate INPs or GMCs. Reprogramming tumor NBs to bypass defective immature INPs and generate GMCs instead halt tumor growth in *brat,patr-1* hypomorphic brains.

### Regulation of cell intrinsic competency to generate transit-amplifying progenitors

The inability to directly assess the capability of radial glia to generate TAP subtypes *in situ* has hindered our understanding of the generation of diverse cell types during neurogenesis. The obstacle mainly stems from lack of insight into cell-type-specific enhancers that are functionally relevant to TAP formation. We leveraged the well-established cell type hierarchy in NB lineages to define a large cohort of active neurogenic enhancers in T2NBs including cell-type-specific enhancers of genes essential for INP formation during fly larval brain neurogenesis (Fig. 1C). Two models for the regulation of NB capability to generate INPs can be deduced from previous studies ^13–17,31–33^. The deterministic model suggests that cell-type-specific enhancers of genes that promote INP formation are inaccessible in T1NBs leading to their inability to generate INPs. This model predicts that inducing T1NBs to generate INPs would necessitate re-establishment of chromatin accessibility at these enhancers (Fig. 1A). By contrast, the hierarchical model suggests that enhancers driving gene expression essential for INP generation are accessible in both NB types but are inactive in T1NBs (Fig. 1A). Our study revealed that these cell-type-specific enhancers display characteristics that are indicative of poised enhancers including accessible chromatin, high H3K27me3/H3K4me1 and low H3K27ac in T1NBs (Fig. 1D,F; S1D,E). These data support the hierarchical model that the capability to generate INPs is inherent to both NBs but activated by lineage-specific transcription factors such as Btd in T2NBs.

What prevents poised enhancers of genes essential for INP formation from activation in T1NBs? Lack of T1NB-intrinsic activators that function analogously to Btd is likely a major factor. This study uncovered a previously unknown function of Ase that instills a barrier mechanism limiting the ability of T1NBs to generate INPs. We identified Pros as a key downstream-effector of Ase-instilled barrier mechanism (Fig. 4; S3). Ase likely limits INP generation by promoting Pros expression, which does not appear necessary for NB function ^31^, suggesting that the barrier mechanism does not directly inhibit activation of poised cell-type-specific enhancers in T1NBs (Fig. 4). Instead, the barrier mechanism likely functions in GMCs to limit reprogramming into immature INPs due their similarity in the transcriptome and genomic landscape (see discussion below). Thus, we propose that Btd and Ase function opposingly in a double-assurance mechanism to ensure the capability to generate INPs is maximally activated in T2NBs while safeguarding this ability in T1NBs.

A perplexing question arose from our study: why are cell-type-specific enhancers of genes essential for INP formation maintained in a poised state in T1NBs despite lacking INP progeny? Genomic studies in embryonic, hematopoietic and skin stem cell lineages have reported the identification of a large cohort of poised enhancers ^46,47,61^. Subsets of these poised enhancers transition to an active state marked by increased H3K27ac and concomitant decrease in H3K27me3. Thus, a function of the poised chromatin state is likely to bookmark developmental enhancers for rapid activation in a response to signaling and developmental cues. Many poised developmental enhancers do not appear to become activated and eventually lose chromatin accessibility in differentiated cell types. These enhancers might serve as a reservoir of *cis*-regulatory elements that are in a low-barrier state allowing stem cells to adapt to environmental changes by readily changing gene expression. Our study suggests that similar poising mechanisms appear to regulate cell-type-specific enhancers of genes in neural differentiation programs in neural stem cells likely to ensure timely and rapid neuronal differentiation.

### Transit-amplifying progenitor plasticity

It is currently unknown what constrains radial glial progeny to assume the identity of a specific TAP subtype and whether their identities can be interconverted. In *Drosophila*, an INP can undergo 5-6 rounds of asymmetric division to generate a GMC each time, but a GMC always divides symmetrically to generate 2 neurons suggesting that these two types of NB progeny exhibit drastically different neurogenic potential. Post-embryonically removing *btd* function in T2NBs has no effects on their lineage-specific marker gene expression and does not detectably upregulate Ase and Pros protein expression. Yet, their presumptive immature INPs assume a GMC fate marked by Pros protein expression ^20^. Similarly, elevating Ase expression in T2NBs does not affect lineage-specific marker expression but leads to their progeny to assume a GMC fate (Fig. 3E). These findings suggest that (1) GMC fate is the default cell identity for NB progeny in both lineages and (2) the expanded neurogenic potential in immature INPs arise from superimposition of the immature INP-specific genetic program upon the GMC identity. Consistently, analyses of published single-cell mRNA sequencing datasets of wild-type larval brains indicate that GMCs in both NB lineages are most closely related to Ase^-^ immature INPs followed by Ase^+^ immature INPs (Fig. 5A). Furthermore, the vast majority of active neurogenic enhancers in T2NBs remain accessible in immature INPs and likely in GMCs (Fig. 1C)^38^. Elevated Pros expression is sufficient to reprogram Ase^-^ immature INPs into GMCs without altering their lineage identity as indicated by their progeny extending bifurcating axon projections that are characteristic of T2 lineage-specific neurons (Fig. 4D,G)^31^. Thus, the identities of GMCs and immature INPs can be interconverted by regulators of NB capability to generate INPs. In agreement with this model, elevated Ase expression in tumor NBs in adult *brat*-mutant brains results in bypassing the generation of differentiation-deficient INPs, which serve of cells-of-origin for tumor cells, and generate GMCs instead halting tumor expansion (Fig. 6, Fig 7B).

Poised enhancers identified in vertebrate stem cell types likely represent regulatory potential of a large repertoire of gene expression programs that can be activated in their TAP progeny ^46,47,61^. Although subset of these enhancers become activated in TAPs to promote lineage-specific differentiation programs, most remain in a poised state and display diminished accessibility much later in development. Similar to *Drosophila*, vertebrate TAPs likely are significantly more plastic than what their classifying subtypes suggest, potentially increasing their susceptibility to tumorigenic transformation and leading to childhood malignancies ^62,63^. Insight into reprogramming the activity of developmentally poised enhancers in tumor cells might allow for the generation of distinct TAP subtypes bypassing differentiation blockades that drive tumor expansion.

### Limitation of this study

Our study suggests that T1NBs maintain cell-type-specific enhancers of genes essential for INP formation in a poised state and can generate immature INPs that differentiate to INPs when induced by elevated Btd expression. The profile of histone modifications at these cell-type-specific enhancers correlates with poised enhancers. Demonstration of the concurrent presence of H3K27me1 and H3K27me3 at these enhancers is required to meet the definition of poised enhancers.^46^ Induced immature INPs generated by T1NBs display cell identity and cell proliferation markers that are consistent with their differentiation to INPs. Our current data do not indicate that these induced INPs are functionally analogous to endogenous INPs.^18^

## Acknowledgement

We would like to thank Drs. M. Harrison, U. Walldorf and D. Yamamoto for sharing antibody reagents. We would like to thank the Bloomington *Drosophila* Stock Center and the Vienna *Drosophila* Resource Center for fly stocks and Developmental Study Hybridoma Bank for antibodies. We would like to thank Genetivision for generating transgenic fly lines. We would like to thank Dr. M. Harrison and members of the Lee lab for comments on the manuscript. This work was supported by a grant from the National Institute of Neurological Disorders and Stroke grants (R01NS134942 to C.-Y.L).

## Materials and Methods

### Fly genetics and transgenes

For experiments driving a *UAS*-transgene and the matched controls, crosses were carried out in 6-oz plastic bottles, and eggs were collected on apple caps in 8-hour intervals. Larvae were kept at 25°C and 24 hours after collection were genotypes and transferred to cornmeal caps. Larvae were shifted to 33°C for 72 hours to induce *UAS*-transgene expression for overexpression or knock down unless otherwise specified. *Tubulin-Gal80*^*ts*^ was used to prevent transgene expression in the embryo for all experiments without heat shock induction. *Worniu-Gal4* was used to drive Pan-NB expression, *31F04-Gal4* (*Tll-Gal4*) was used to drive T2NB specific expression, and *9D11-Gal4* on the second chromosome (*Erm-Gal4*) was used to drive immature INP expression. Experiments shown in Fig. 2A and Fig. 4L-N utilized the same number of *UAS*-transgenes between control and experimental to account for driver dilution.

GFP reporter transgenes used were cloned with Gateway LR Clonase (ThermoFisher Scientific, Waltham, MA) to insert PCR-amplified genomic regions into the VanGlow vector with the DSCP (Addgene#83338). Coordinates and validated sequences for the enhancers used are included in Table S5. For *pros*^*ΔAse*^*-GFP*, gBlocks (Integrated DNA Technologies, Coralville, IA) were ordered with Ase binding sites (CAGCTG) deleted. The reporters were integrated into the VK22 site on chromosome 2 using φC31-mediated integration (Genetivision Inc). Transgenic flies were identified in the F1 generation based on their red eye color. Expression was validated by dissecting and imaging wandering third instar larvae.

For generation of *UAS-btd::3XMyc*, open reading frame for *btd-RA* with a 3XMyc epitope added, was cloned into the *pUASt-attB* vector through use of standard PCR, restriction digest and ligation procedures. The Transgene were inserted into the *pUAST-attB M(3xP3-RFP*.*attP)ZH-86Fb* docking site using ΦC31 integrase-mediated transgenesis.^64^ DNA injections were carried out by Genetivision Inc.

### Immunofluorescent staining and antibodies

Larval brains were dissected in phosphate buffered saline (1X PBS) before fixation for 23 minutes at 25°C in a solution containing 100 mM PIPES (pH 6.9), 1 mM EGTA, 0.3% Triton X-100 and 1 mM MgSO4 and 4% formaldehyde. Samples were then washed in a solution 1X PBS and 0.3% Triton X-100 (1X PBST). Samples were incubated in a solution of 1X PBST and primary antibody (found in Key Resources Table) for 3 hours at. Samples were washed with 1X PBST before incubated overnight at 4°C in a solution of 1X PBST and secondary antibodies or phalloidin (found in Key Resources Table). The following day samples were washed in 1X PBST before equilibrating in ProLong Gold antifade mount (ThermoFisher Scientific). Adult brain samples were prepared using the same protocol with the adjustment of performing both primary and secondary antibody incubation steps overnight at 4°C.

### Hybridization Chain Reaction (HCR) and immunofluorescent staining

Experiments where mRNA transcripts in larval brains were quantified, was done by performing in situ HCR v3.087. All cDNA sequences utilized NCBI to generate HCR probes. HCR probes can be found in the Key Resources Table. Samples were dissected in 1X PBS and fixed for 23 minutes in a solution of 100 mM PIPES (pH 6.9), 1 mM EGTA, 0.3% Triton X-100 and 1 mM MgSO4 with 4% formaldehyde concentration. After fixing, samples were washed in a solution 1X PBS and 0.3% Triton X-100 (1X PBST). Samples were then incubated in a hybridization buffer (10% formamide, 5×SSC, 0.3% Triton X-100 and 10% dextran sulfate) at 37°C for 1 hr. Following the incubation, 5 nM of primary mRNA HCR probes (Molecular Instruments, Los Angeles, CA) were added to each sample and incubated overnight at 37°C. The following day, samples were washed with SSC buffer (10% formamide, 5×SSC, 0.3% Triton X-100) and were incubated for 30 minutes at 25°C with incubated with amplification buffer (5×SSC, 0.3% Triton X-100 and 10% dextran sulfate). Secondary hairpins (Molecular Instruments, Los Angeles, CA) were denatured at 95°C for 2 minutes and incubated on ice for 5 minutes before 3 µM of each hairpin were added to each sample. Samples were incubated overnight at 25°C. The following day samples were washed in 1X PBST before fixing for 12 minutes to initiate immunofluorescent staining protocol as listed above.

### Edu Labeling

Larvae for Edu labeling were collected in 6-oz plastic bottles, and eggs were collected on apple caps in 8-hour intervals. Newly hatched larvae were genotyped and allowed to grow on corn meal caps at 25°C before shifting to 33°C for 48 hours to induce *UAS-ase* expression. This was to ensure the larvae would not be too old to take up Edu containing food. Kankel-White medium caps were prepared with bromophenol blue and with Edu concentration 0.2 mM as previously described.^65^ Feeding was carried out for three hours. Larvae that took up the Edu containing food as indicated by blue food in the guts were dissected and processed for antibody staining as described above. After staining, samples were treated with Click-iT™ EdU Cell Proliferation Kit for Imaging, Alexa Fluor™ 555 dye (ThermoFisher Scientific) as per manufacturer instructions. Following this, samples were washed in 1X PBST before equilibrating in ProLong Gold antifade mount (ThermoFisher Scientific).

### Imaging and Quantification of immunofluorescent samples

Images of samples were acquired on a Leica SP5 scanning confocal microscope (Leica Microsystems Inc) using a 63X glycerol immersion objective. Images were taken at 1.5X zoom for whole lobe, or 3X zoom for single neuroblasts or clonal lineages. Whole brain lobe images were taken with z-stacks of 1.51 µm thickness, while 3X zoom images and all smFISH images used 0.5 µm thickness.

Quantification was done where the observer was blind to the genotypes. All statistical analyses were performed using a two-tailed Student’s t-test with p-values <0.05 (*), <0.005(**), <0.0005 (***), and <0.00005 (****). GraphPad Prism was used to generate dot-plots. Only the central brain was looked at and identified based on morphology. Type I and type II NBs were identified based on size (>7 µm) and expected marker profile or labeling with mcd8:GFP or mcd8:RFP. INPs were identified by small size (<5 µm) and were Dpn^+^Ase^+^. Experiments targeting T1NBs in Fig 2 used the ventral side of the brain where only T1NBs are present.

The number of INPs was manually counted using only one brain lobe per brain. Fractions of T2NBs were calculated by counting all T2NBs in one brain lobe per brain and counting the number of T2NBs that met criteria outlined in each experiment. Fraction of T1NBs generating immature INPs was calculated by counting all T1NBs on the ventral side of the brain and counting the number of T1NBs that had Myc^+^Erm^+^Dpn^-^ small cells directly adjacent. Quantification of foci was done by manually counting the number of foci found in the T2NB in every other z-stack and normalizing to mRpl15 foci. Only one NB per brain was quantified to ensure biological variability. For pixel intensity comparisons, the pixel intensities of Dpn or Ase were measured in nucleus of cells of interest by using Image J software and selecting an ROI and the pixel intensities of GFP reporter in the identical area were measured.

### Collection of brains for ATAC-seq, CUT&RUN, and bulk RNA-seq

T2NB enriched brains were acquired as previously described ^24,38^ where *brat*^*11/Df*^ brains were collected in 8 hour windows incubated for 120 hours at 25°C. Immature INP and INP enriched brains were acquired as described where pan-NB driver WorGal4 was used to drive *UAS-insb* in *brat*^*11/Df*^ brains to induce synchronous differentiation by using *Tub-Gal80*^*ts*^ to control when Insb expression is activated. ^24,38^ Samples were collected in 8-hour windows and kept at 25°C for 96 hours before shifting to 33°C to induce Insb expression. All INP labeled samples utilized a 24-hour induction. T1NB enriched brains utilized a T1NB specific driver *Ase-Gal4* to mis-express *UAS-aPKC*^*caax*^ to prevent T1NBs from differentiating as previously described.^66^ These samples were collected in 8-hour windows and were shifted to 33°C for 72 hours after larval hatching. All samples were dissected in 1X PBS before further processing.

### ATAC-seq on NB or INP enriched brains

ATAC-seq was performed as described in ^67^. Three replicates were collected with at least 5 brains per sample. Samples were collected in 1X PBS and spun down for 3 minutes at 600 g. Supernatant was removed and the brains were resuspended in 50 µl lysis buffer (10 mM Tris pH 7.5, 10 mM NaCl, 3 mM MgCl2, 0.1% NP-40). Samples were transferred to a dounce homogenizer and were dounced at least 40 times on ice. Dounce was washed with 50 µl lysis buffer. 100 µl of sample was transferred to a PCR tube and spun for ten minutes at 500 g at 4°C. Supernatant was removed and pellet was resuspended in a mix containing 5 µl Transposase Buffer, 2.5 µl sterile water, and 2.5 µl Tn5 transposase all sourced from Tagment Illumina. Samples were incubated at 37°C for 30 minutes. DNA was then purified using Qiagen MiniElute Reaction Cleanup Kit (Qiagen) and eluted in 10 µl EB Buffer. DNA was combined with 10 µl Illumina Index, 5 µl water, and 25 µl NEB Next Hi-Fi 2x PCR master mix. DNA was then PCR amplified 12 cycles using the following conditions: 72°C for 5 minutes, 98°C for 30 seconds and then followed by 12 cycles of 98°C for 10 seconds, 63°C for 30 seconds and 72°C for 1 minute. Libraries were purified using 1.1 x AMPure Beads without size selection. (Beckman Coulter). Sequencing utilized IlluminaNovaSeq-6000 chip (University of Michigan Advanced Genomics Core) with 150-bp paired-end reads.

### CUT&RUN on NB enriched brains

Samples were prepared as previously described.^67,68^ Brains were dissected in under 45 minutes with at least 50 brains per replicate for TFs and 25 brains per replicate for histone marks. Two replicates were collected for each sample.. Larval brains were dissected in 1X PBS, transferred to 0.5mL Eppendorf tubes, and spun down for 3 minutes at 600 g and supernatant was removed. Samples were then prepared using CUTANA ChIC/CUT&RUN (Epicypher) per the manufacturer’s protocol with some modifications. Brains were resuspended in 100 µl wash buffer before being transferred to a dounce homogenizer and were dounced at least 40 times on ice. Sample was transferred to a 1.5 ml tube and spun for 3 minutes at 600 g. Pellet was processed using CUT&RUN kit. For each sample, 0.5 µg of antibody was used (or 0.5 µl if antibody concentration was unknown). Additionally, 0.5 ng of *Escherichia coli* spike-in DNA was added into every sample as a spike-in control.

Samples targeting acetylation utilized 10 mM of sodium butyrate (Sigma Aldrich) in all buffers. Samples using mouse antibodies were processed with an additional antibody incubation step, in which samples were washed twice with cell permeabilization buffer after primary antibody incubation, and then were incubated for 1 hour with 0.5 µg of rabbit anti-mouse IgG (Abcam, ab46540).

DNA was diluted to 50 µl in 0.1x TE which was followed by library prep using the NEBNext Ultra II DNA Library Prep Kit for Illumina (E7645) along with NEBNext Multiplex Oligos for Illumina (E6440). The manufacturer’s protocol was followed with modifications. Adaptor was diluted by a ratio 1:25. The bead clean-up steps were performed using 1.1 x AMPure Beads without size selection. The PCR cycle was adjusted to match the settings listed in CUTANA ChIC/CUT&RUN kit. DNA was eluted in 20 µL 0.1 x TE. Agilent TapeStation was used to assess DNA quality and concentration. If samples had greater than 1% adaptor, additional bead cleanup was done. Samples were sequenced on IlluminaNovaSeq-6000 chip (University of Michigan Advanced Genomics Core) with 151-bp end reads.

### RNA-seq on NB or INP enriched brains

Samples were prepared as previously described.^24^ Samples were dissected in 20-minute intervals and at least 50 brains were collected. All samples were prepared in triplicate. Samples were prepared in RNase-free conditions. RNA was isolated using TRIzol (ThermoFisher Scientific) and mRNA was then purified using the RNeasy Micro Kit (Qiagen) according to the manufacturer’s instructions. Libraries were prepped using SMART-Seq Stranded Kit (Takara, 634444) according to manufacturer’s instructions. RNA was sequenced on IlluminaNovaSeq-6000 chip with 150bp end paired reads (University of Michigan Advanced Genomics Core).

### Re-analysis of scRNA-seq data

Processed scRNA-seq data from L3 brains consisting of all NB lineages^35^, T2 lineage^36^, and INPs^34^ were downloaded from NIH GEO. Datasets were merged, and QC filtering was performed using scanpy^69^ (min genes per cell = 200, min cells per gene = 3, mitochondrial percent < 5). Due to batch effect across datasets, we re-annotated the merged dataset using known marker genes (T2NB: *fru*^*+*^, *pnt*^*+*^, *dpn*^*+*^, *erm*^*-*^, *ase*^*-*^; Ase^-^ immature INP: *fru*^*-*^, *pnt*^*+*^, *erm*^*+*^, *ase*^*-*^, *dpn*^*-*^, *opa*^*-*^, *dap*^*-*^; Ase^+^ immature INP: *fru*^*-*^, *pnt*^*+*^, *erm*^*+*^, *dpn*^*-*^, *opa*^*-*^, *dap*^*-*^; INP: *dpn*^*+*^, *ase*^*+*^, *fru*^-^, *opa*^*+*^, *dap*^*-*^; T1NB: *fru*^*+*^, *ase*^*+*^, *dpn*^*+*^, *pnt*^*-*^, *erm*^*-*^; T1 GMC: *dpn*^*-*^, *ase*^*+*^, *Sp1*^*-*^, *opa*^*+*^; T2 GMC: *dpn*^*-*^, *ase*^*+*^, *Sp1*^*+*^, *opa*^*+*^; immature neurons: *Hey*^*+*^, *fru*^*-*^, *ase*^*-*^; neurons: *nSyb*^*+*^, *ase*^*-*^). Known marker genes were visualized by cell-type using scanpy dotplot.

Gene were declare on or off based on whether they were expressed above average (z-score > 0) within a given cell-type. Overlapping and differential genes were visualized using matplotlib venn3 [doi: 10.1109/MCSE.2007.55]. Transcription factors were annotated based on FlyBase’s (release FB2024_04) transcription factor category.

Cell type similarity was assessed in two ways. Dendogram based on pearson correlation between cell-types was generated using scanpy dendrogram function. Eucledian distance of the average gene expression between cell-types was caulculated using scipy pdist module ^70^, and visualized using seaborn heatmap [doi: https://doi.org/10.21105/joss.03021.].

### Data processing and analysis of ATAC-seq

ATAC-seq datasets for T2 and INP enriched brains^38^ as well as T1 enriched brains were processed. Quality control was performed on FASTQ files using FastQC and FastQ-Screen (https://www.bioinformatics.babraham.ac.uk/projects/fastqc/).^71^ Illumina adapters were trimmed from FASTQ files using NGmerge.^72^ Trimmed reads were aligned to the dm6 genome using bowtie2 using flags: --very-sensitive -I 10 -X 2000.^73^ Duplicate reads were removed using samblaster^74^, and data quality was checked using ATAQV^75^. Preliminary bigwig files were generated using deeptools bamCoverage.^76^ MACS2 was used to call peaks using default parameters.^77^

TMM (Trimmed Mean of M-Values) normalization was performed to accout for difference in library composition and sequencing depth. First, all peaks from MACS2 were merged, and blacklisted regions determined by ENCODE were removed.^78^ Then, reads in peaks were counted using featureCounts on each individual replicate.^79^ EdgeR calcNormFactors (method = TMM) was colled on each count matrix, and the normalization factor was calculated as 1,000,000/(normFactor * number of reads in peaks).^80^ Bigwig files were generated for individual replicates using deeptools bamCoverage on aligned bam files with the scaleFactor equal to the final normalization factor that was calculated, and only mono/sub nucleosomal fragments were kept using maxFragmentSize 120. Bigwigs were then merged using wiggletools write_bg^81^, converted back to bigwig files using bedGraphToBigWig^82^ and z score bigwig files for histone marks were generated and visualized in IGV^83^.

### Data processing and analysis of CUT&RUN

CUT&RUN datasets for T2 enriched brains as well as T1 enriched brains were processed.^36,38^ Quality control was performed on FASTQ files using FastQC and FastQ-Screen. Illumina adapters were trimmed from FASTQ files using NGmerge. Trimmed reads were aligned to the dm6 genome using bowtie2 using flags: --very-sensitive -I 10 -X 2000. Duplicate reads were removed using samblaster. Preliminary bigwig files were generated using deeptools bamCoverage. Transcription factor peaks were called following the CUT&RUNTools2 pipeline^84^, blacklist regions were removed, and the remaining peaks were merged using bedtools.

TMM (Trimmed Mean of M-Values) normalization was performed to accout for difference in library composition and sequencing depth for bigwig files. First, all peaks from each histone mark were called using gopeaks^85^, broad peak calling was used for H3K27me3 and H3K27ac, and narrow peak calling was used for H3K4me1 and H3K4me3, Peaks for each type of histone mark were merged, and blacklisted regions determined by ENCODE were removed. For transcription factors, peaks generated from CUT&RUNTools2 were used. Reads in peaks were counted using featureCounts on each individual replicate. EdgeR calcNormFactors (method = TMM) was colled on each count matrix, and the normalization factor was calculated as 1,000,000/(normFactor * number of reads in peaks). Bigwig files were generated for individual replicates using deeptools bamCoverage on aligned bam files with the -- scaleFactor equal to the final normalization factor that was calculated. Bigwigs were then merged using deeptools bigwigCompare, and z score bigwig files were generated and visualized in IGV.

### Data processing and analysis of RNA-seq data

Bulk RNA-seq datasets for T1, T2, immINP, and INP enriched brains were processed. Quality control was performed on FASTQ files using FastQC and FastQ-Screen. Illumina adapters were trimmed from FASTQ files using cutadapt [doi: https://doi.org/10.14806/ej.17.1.200]. Trimmed reads were aligned to the dm6 transcriptome using hisat2 using flags: --no-mixed –no-discordant.^86^ A gene by count matrix was generated using featureCounts to count alignments from the hisat2 output to the annotated dm6 transcriptome (refGene). The resulting matrix was processed using DESEQ2 ^87^, and a z-score scaled heatmap of known marker genes was generated from the variance stabilized transformed matrix.

### Flowcharts, heatmap, and gene list analyses

To determine regions that fulfilled designed criteria, we used bedtools to filter regions based on overlapping peak criteria.^88^ ATAC-seq peaks called by MACS2 were interesected across all replicates (keeping only peaks that are called in every replicate), merged, and ENCODE blacklist regions were removed. Promoters were determined as 1000bp upstream of TSS, and regions overlapping with promoters were also removed. For all analyses, we started from the set of accessible regions that are open in type II neuroblasts, as T2NB accessibility is a superset of all accessible regions by visual inspection. Transcription factor peaks were called following the CUT&RUNTools2 pipeline^84^, merged using bedtools, and ENCODE blacklist regions were removed. Peaks were associated with their nearest gene using homer annotatePeaks^89^, and genes lists from the scRNA-seq data were used to associate regions with cell-types that were enriched for the associated gene. Some peaks were manually reassigned to their nearest gene, in the case where Homer misannotates due to the presence of ncRNAs. These cases are highlighted in table S1. Data was integrated using pandas [10.25080/Majora-92bf1922-00a] in python, and resulting tables are included within the supplemental data.

We performed k-means clustering on the H3K4me1, H3K27ac, and H3K27me3 CUT&RUN data from type I neuroblast enriched brains using deeptools computeMatrix and plotHeatmap. We performed k-means clustering with n=3 clusters on the set of regions defined by as being accessible in type II neuroblasts, not including promoters, and used the z-scored bigwig files for each histone mark. Clusters were extracted from the matrix using plotHeatmap, named based on the histone mark pattern (Active: H3K4me1^+^, H3K27ac^+^; Poised: H3K4me1^+^, H3K27me3^+^; Inactive: H3K4me1^-^, H3K27ac^-^), and interesected using pandas in python as well as visualized.

## Supplemental Figure Legends

**Supplemental Figure 1.**
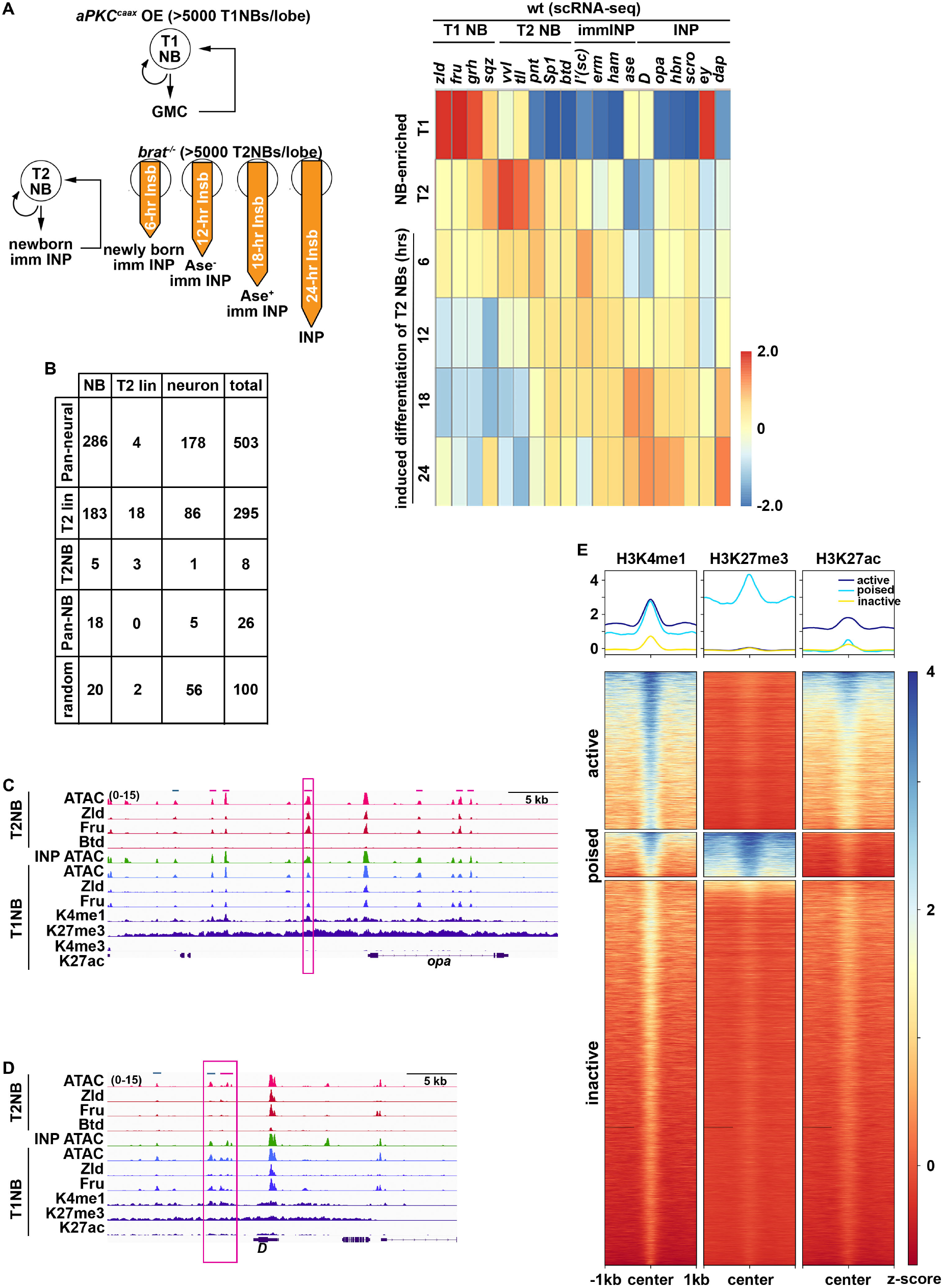
Identification of active neurogenic enhancers in fly larval brain NBs. (A) Schematic for enrichment of NBs and INPs used for isolating Bulk-NB or INP lysate. The distribution of representative gene transcripts in cell types of T1 and T2 lineages derived from single cell RNA-seq of wild-type larval brains shows transcripts abundance consistent with the enriched NB or INP system. (B) Table summarizing the activity pattern of enhancers determined by the flowchart in Fig. 1C. We assessed enhancer activity patterns using lab-generated reporters or against an existing database to validate our approach.^90^ Random refers to randomly selected non-promoter enhancers whose activity pattern was available from the Janelia Flylight website. (C-D) Genome browser tracks showing patterns of chromatin accessibility, TF occupancy and histone marks in indicated cell types in *opa* and *D* loci where the magenta bars correspond to pan-neural enhancers, and teal bars correspond to T2-lin enhancers. Magenta box designates enhancer with INP-specific activity. (E) Heatmap of k-means clustering of H3K4me1, H3K27ac, and H3K27me3 in regions of accessible enhancers in T2 NBs (n=21212, Fig. 1C). Active, poised, and inactive labels are determined based on pattern of histone marks withing cluster.

**Supplemental Figure 2.**
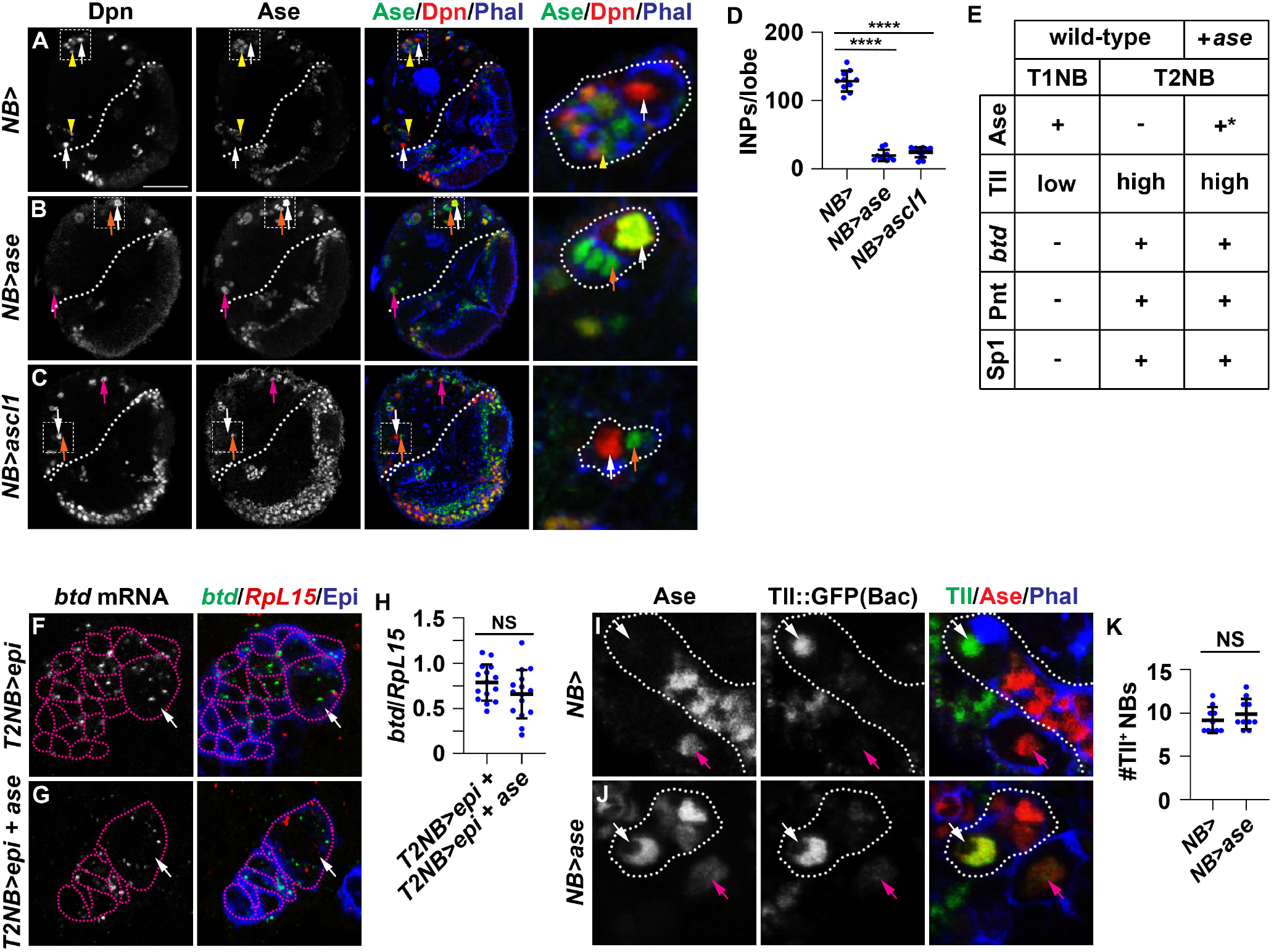
Elevated Ase expression promotes T2NBs to generate GMCs via bypassing INPs. (A-C) Images showing that mis-expressing Ase and Ascl1 with a pan-NB driver (*Wor-Gal4*) results in loss of INPs (small Ase^+^Dpn^+^ cells). Scale bar: 50 µm (D) Quantification showing significant decrease in the number of INPs upon mis-expression of Ase or Ascl1 in T2NBs. (E) Table describing the marker profile of wild-type T1 and T2NBs compared to marker profile after Ase mis-expression in T2NBs. (F-G) Images showing that mis-expressing Ase with a T2NB driver (*Tll-Gal4*) does not alter *btd* transcript abundance. (H) Quantification showing no significant change in the ratio of *btd* vs *Rpl15* transcripts in T2NBs with and without Ase mis-expression. (I-J) Images showing T2NBs mis-expressing Ase using a pan-NB driver (*Wor-Gal4*) still maintain T2NB-specific marker Tll expression. (K) The number of NBs demonstrating high Tll expression is not significantly different between control versus Ase mis-expression. White dotted line circles NB lineage. White dotted line demarcates optic lobe boundary. Magenta dotted line marks the T2NB lineage. White arrow: T2NB. Yellow arrowhead: INP. Magenta arrow: T1NB. Orange arrow: GMC. Scale bar: 10 µm. P-value; NS: Not significant, ****<.00005.

**Supplemental Figure 3.**
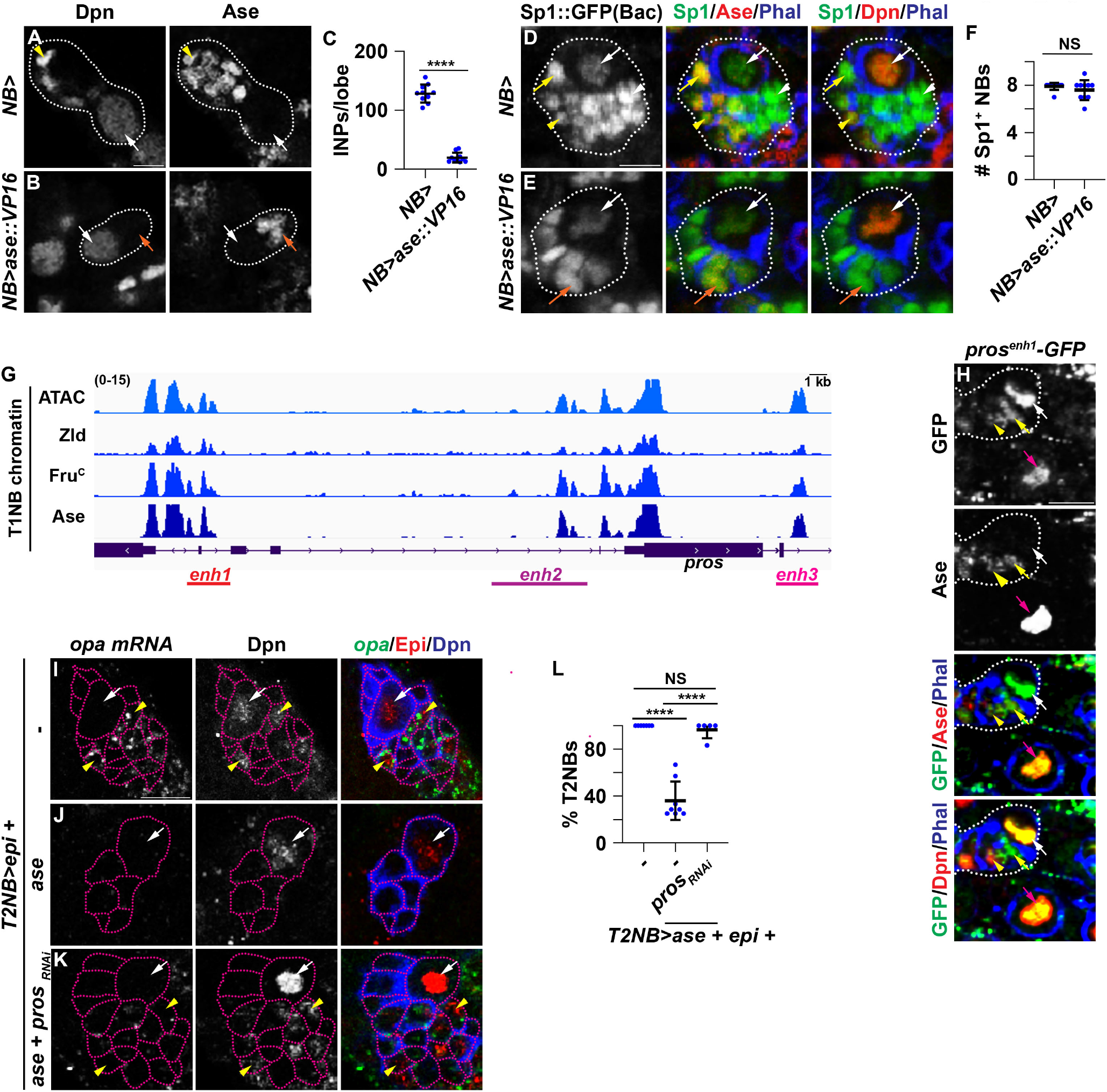
Pros mediates Ase-induced bypass of INP generation in the T2NB lineage. (A,B) Images showing T2NBs mis-expressing a transcriptional activator form of Ase (Ase::Vp16) with pan-NB driver (*Wor-Gal4*) are surrounded by GMCs (small Dpn^-^Ase^+^ cells) while control T2NBs are surrounded by immature INPs (small Dpn^-^Ase^+^ cells) and INPs (small Dpn^+^Ase^+^ cells). (C) Quantification showing that larval brains that mis-express Ase::VP16 in NBs contains drastically fewer INPs per lobe than control brains. (D,E) T2NBs mis-expressing Ase::VP16 with pan-NB driver still maintain T2NB lineage-specific maker Sp1 expression. (F) Upon Ase mis-expression the number T2NBs that display nuclear Sp1::GFP expression is statistically indistinguishable from control T2NBs. (G) Genome browser track of the *pros* locus with patterns of chromatin accessibility and TF occupancy showing three Pan-neural enhancers that are bound by Ase. (H) The expression of a GFP reporter controlled by enhancer 1 from the *pros* locus (*pros*^*enh1*^-GFP) is expressed in T1NBs, T2NBs and INPs. (I-K) Knocking down *pros* function in Ase mis-expressing T2NBs restores their ability to generate INPs. INPs are identified by the expression of *opa* mRNAs and Dpn protein. (L) The percentage of Ase mis-expressing T2NBs generating INPs is restored to a level indistinguishable from wild-type T2NBs following *pros* knockdown. Ase mis-expressing T2NBs are surrounded by drastically fewer INPs than wild-type T2NBs but knocking down *pros* function in Ase mis-expressing T2NBs rescues this defect. White dotted line circles NB lineage. Magenta dotted line marks the T2NB lineage. White arrow: T2NB. Yellow arrow: Ase^+^ immature INP. Yellow arrowhead: INP. Magenta arrow: T1NB. Orange arrow: GMC. Scale bar: 10 µm. P-value; NS: Not significant, ****<.00005.

**Supplemental Figure 4.**
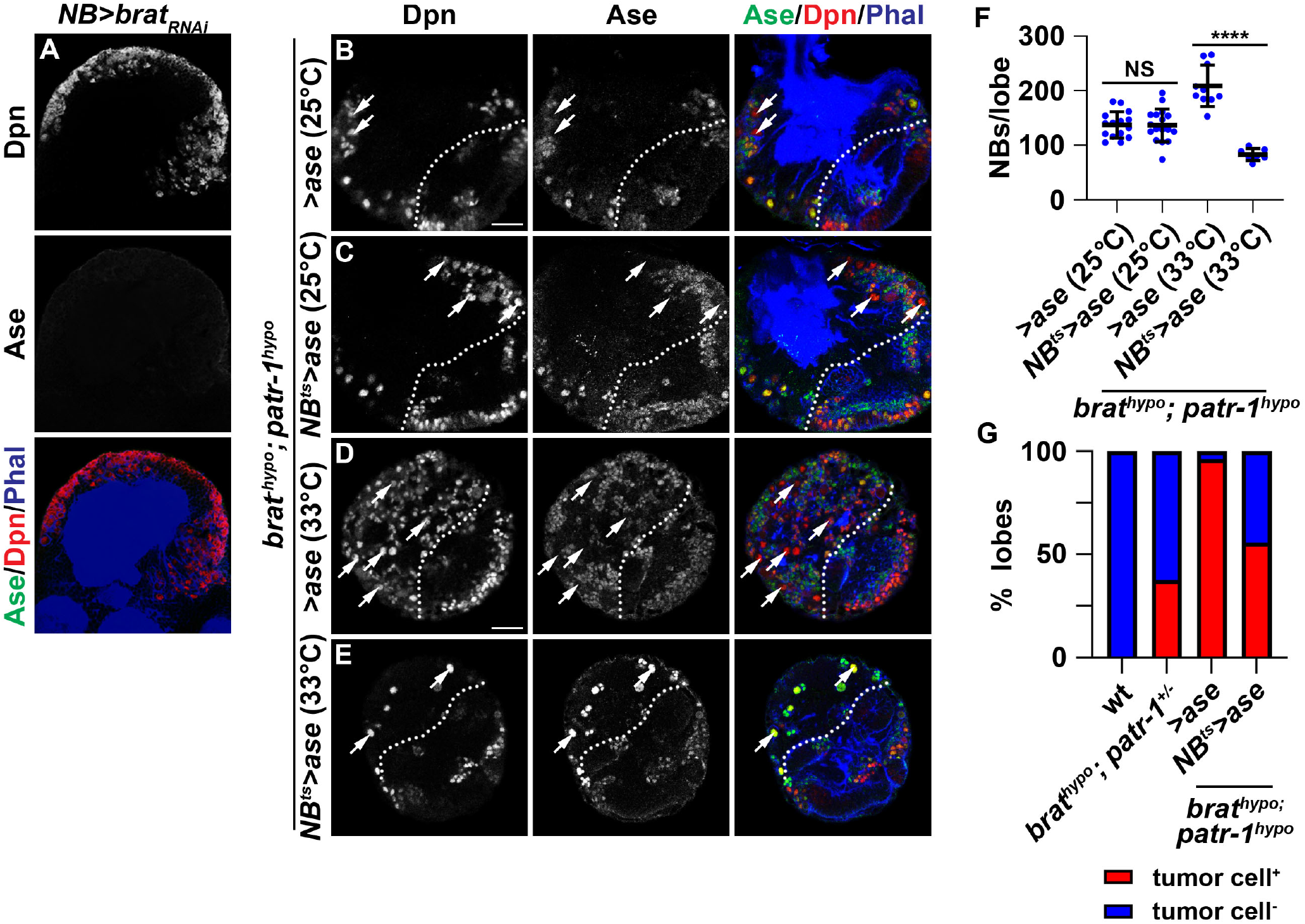
Elevating Ase expression suppresses premalignant NBs formation in *brat,patr-1*-hypomorphic brains. (A) Images showing brat knockdown since larval stage produces T2NBs (Ase^-^Dpn^+^) that persisted into the adult brain. (B-E) Images showing that *brat*^*hypo*^,*patr-1*^*hypo*^ compound larval brains with and without NB-gal4 (*Dpn-Gal4*) when kept at 25°C do not have a significantly different number of NBs, while induction at 33°C will drive Ase mis-expression in the presence of NB-Gal4. Ase mis-expression is sufficient to result in decreased number of T2NBs. (F) Quantification showing that keeping *brat*^*hypo*^,*patr-1*^*hypo*^ compound brain samples at 25°C was able to keep Ase suppressed and with no significant change in the number of NBs. Shifting to 33°C was able to induce Ase mis-expression resulted in a significant decrease in the number of NBs. (G) Quantification showing the fraction of brain lobes where tumors (Dpn^+^) could be detected from between wild-type, *brat*^*hypo*^,*patr-1*^*+/-*^, *brat*^*hypo*^,*patr-1*^*hypo*^, and *brat*^*hypo*^,*patr-1*^*hypo*^ with NB-Gal4 driving Ase samples.

## STAR METHODS

### Key resources table

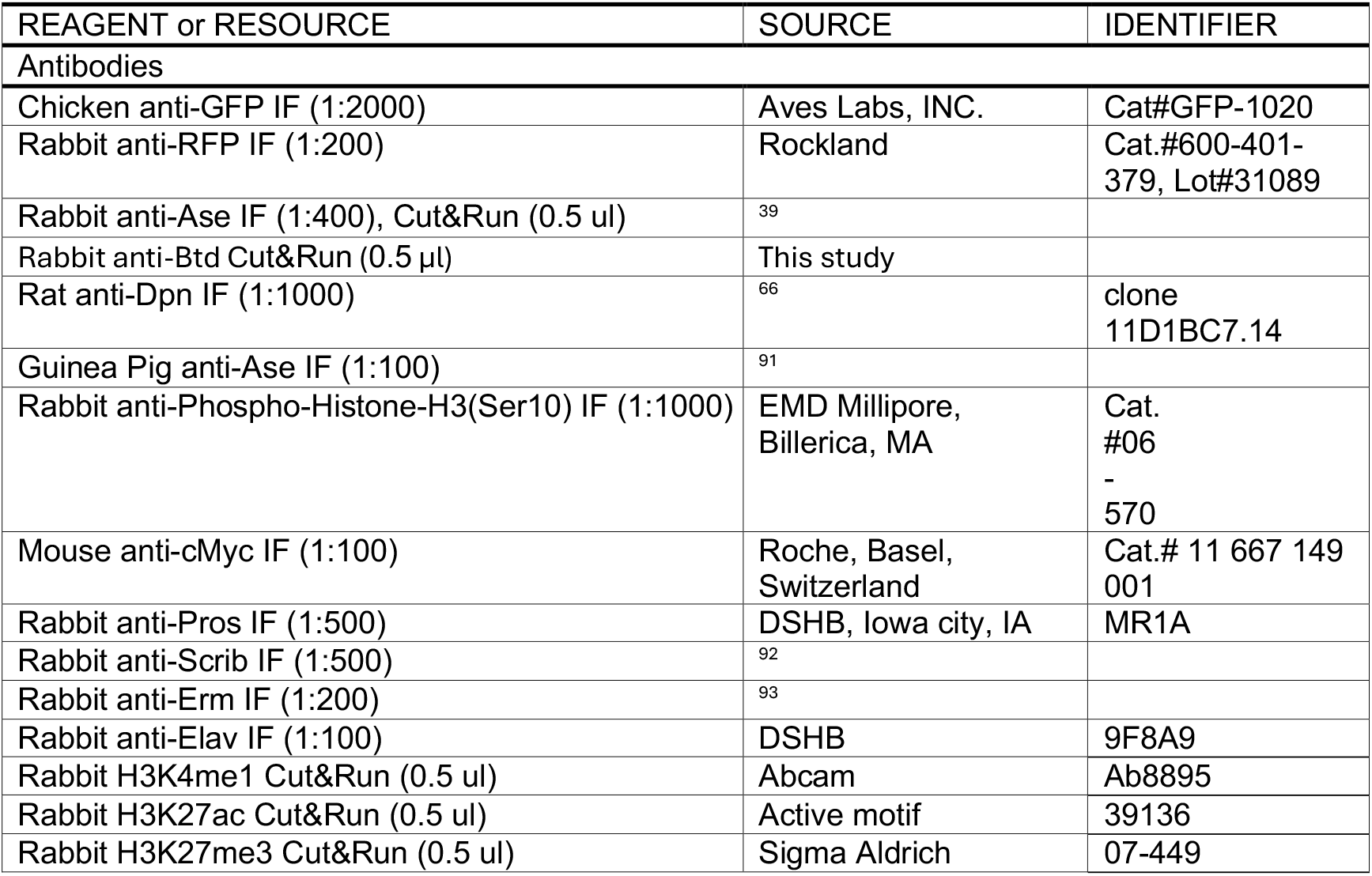
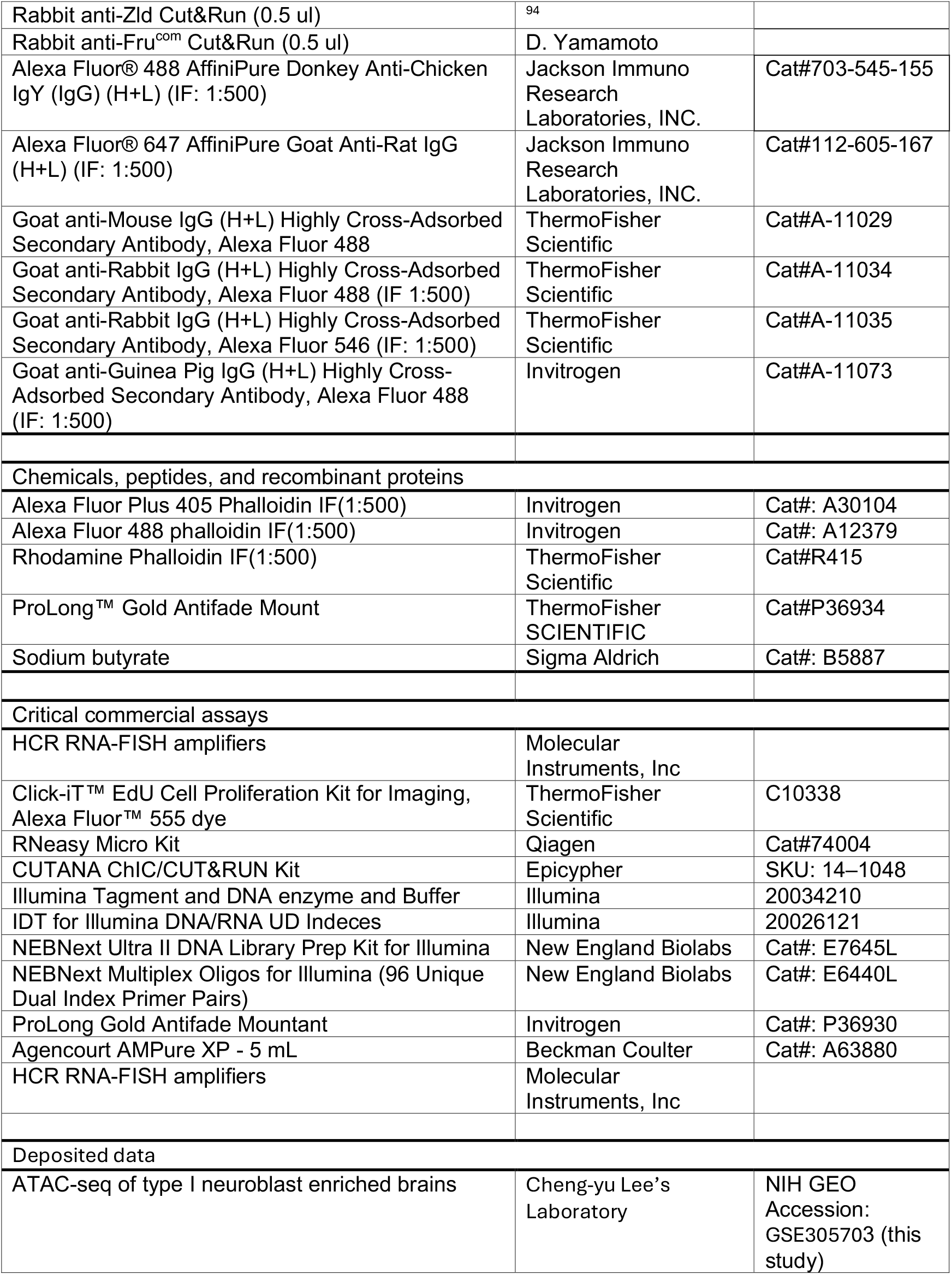
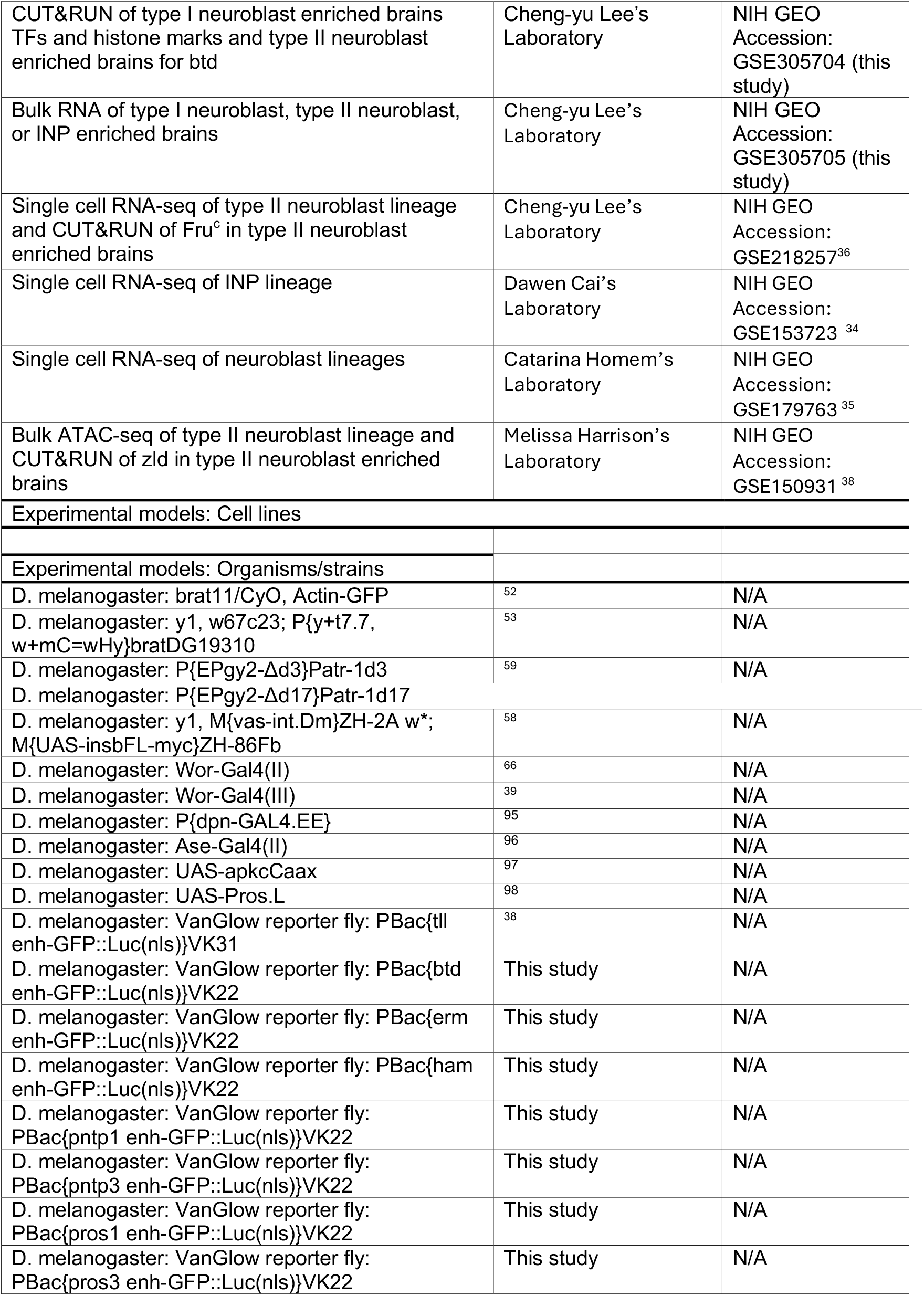
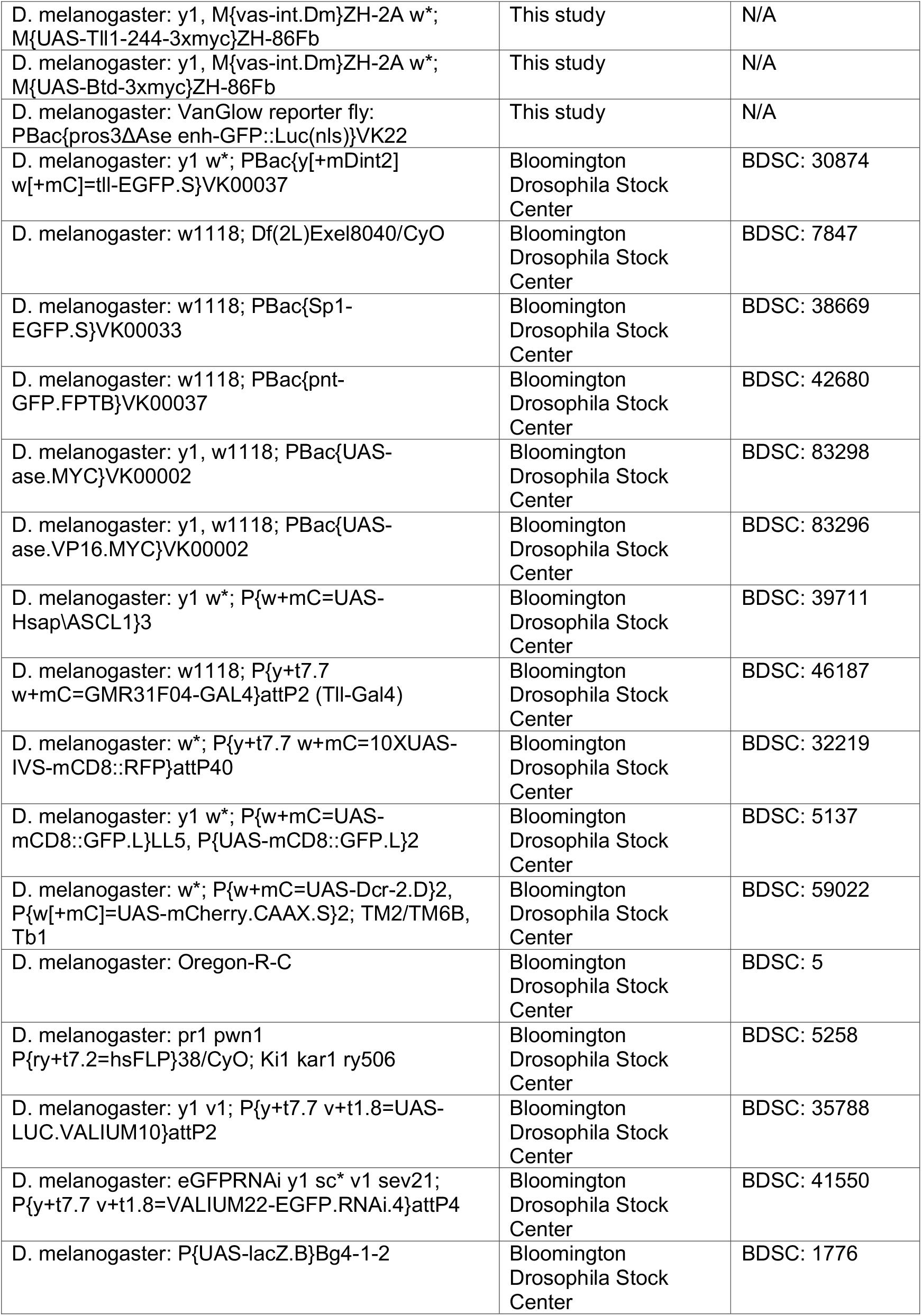
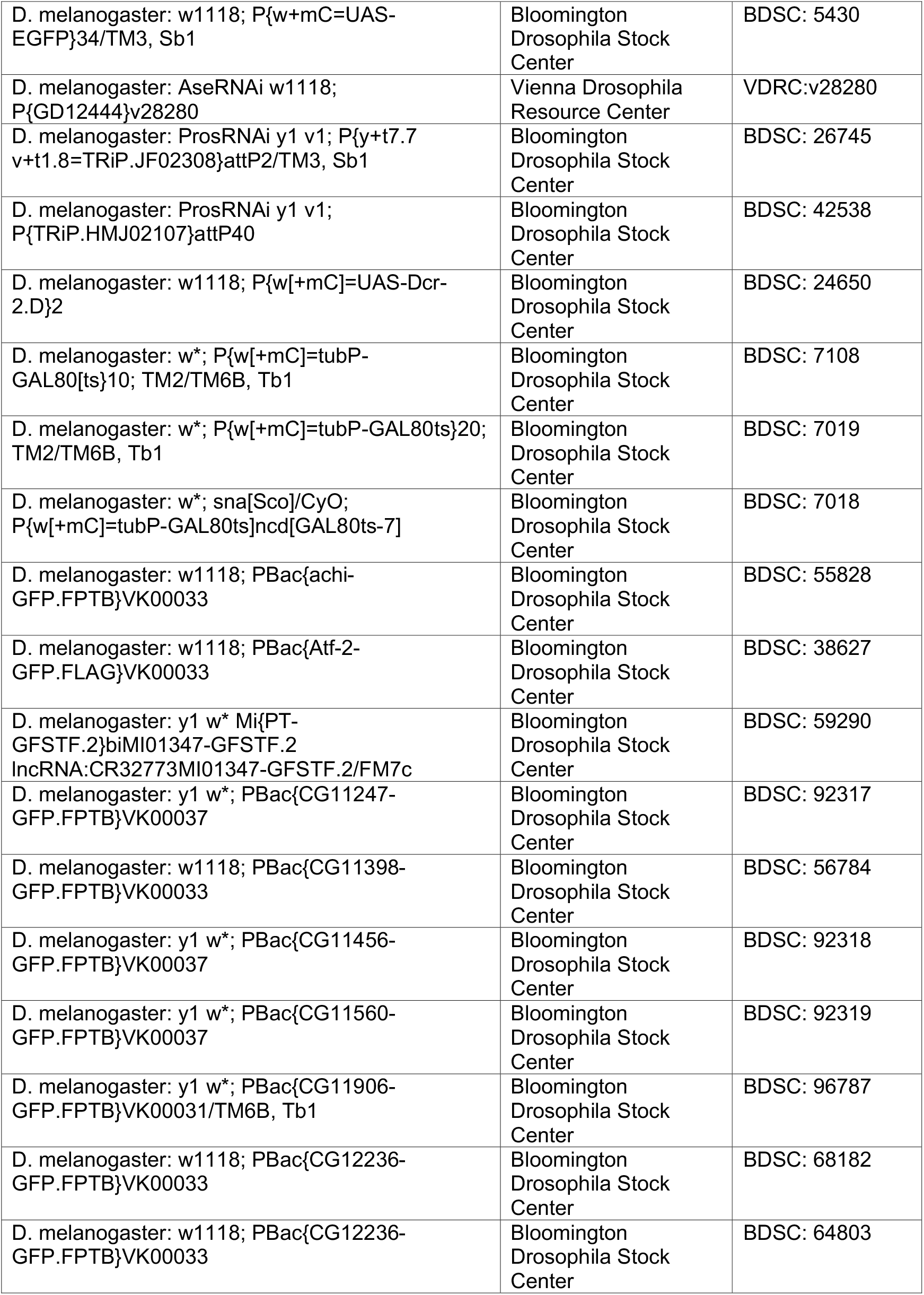
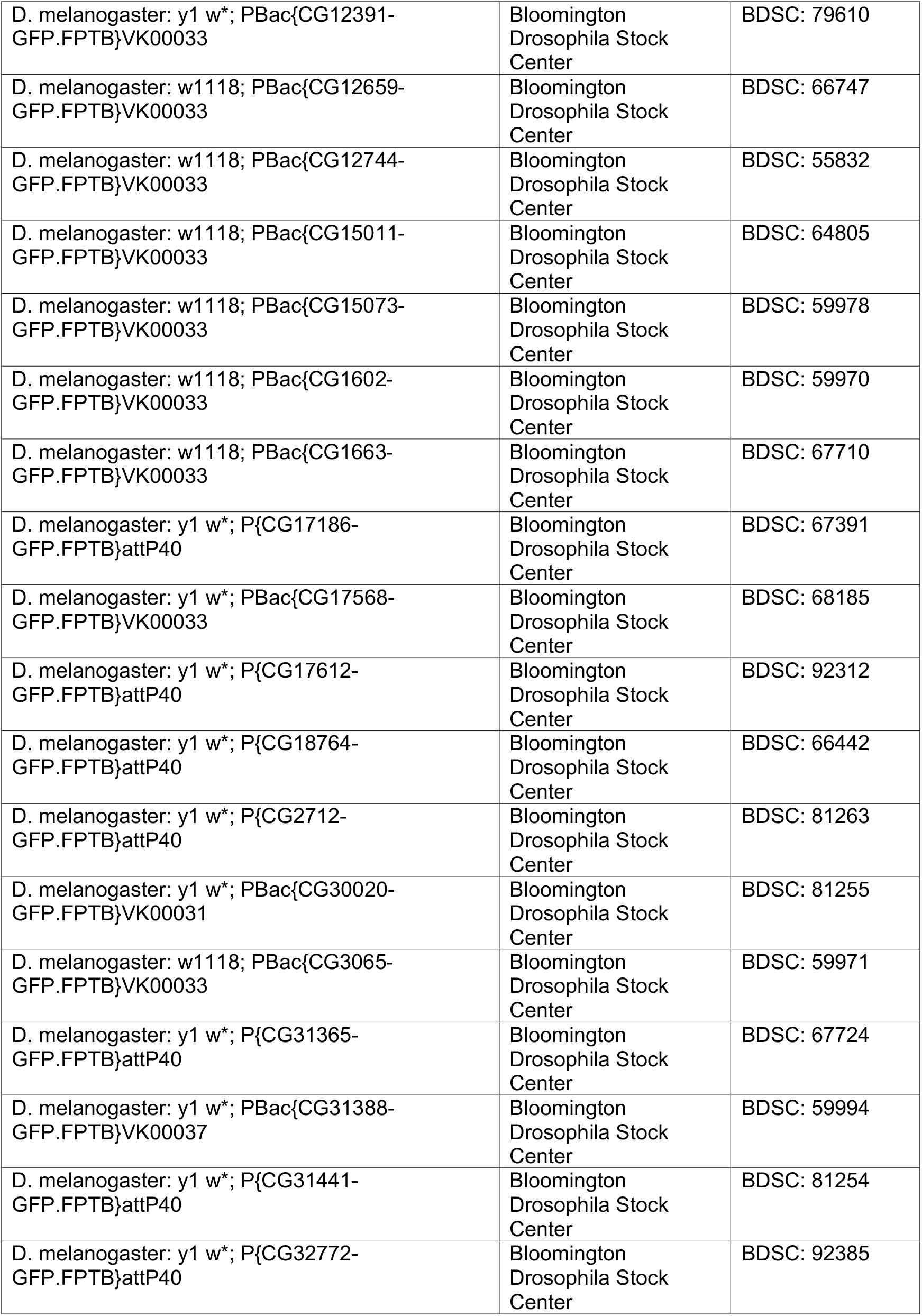
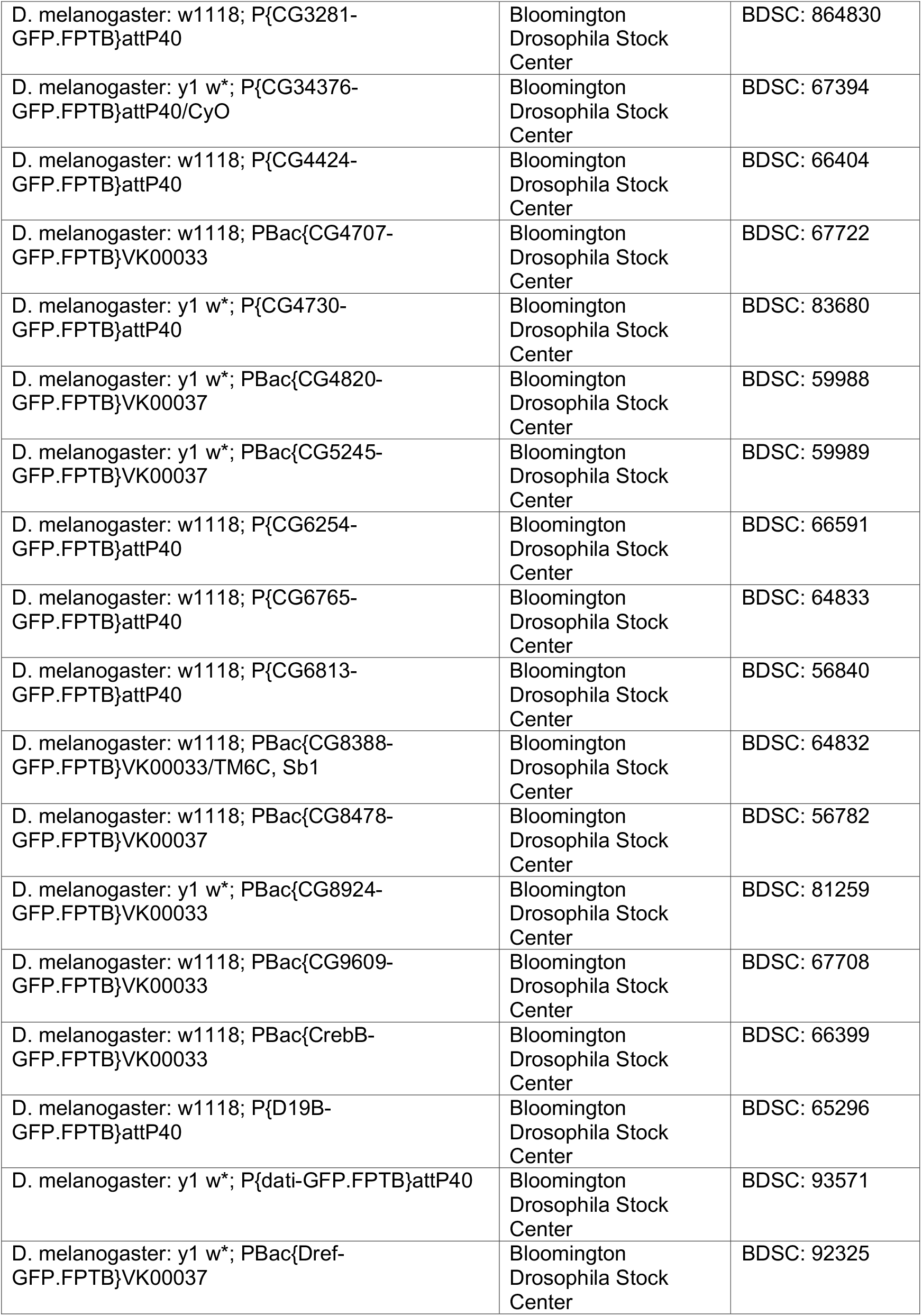
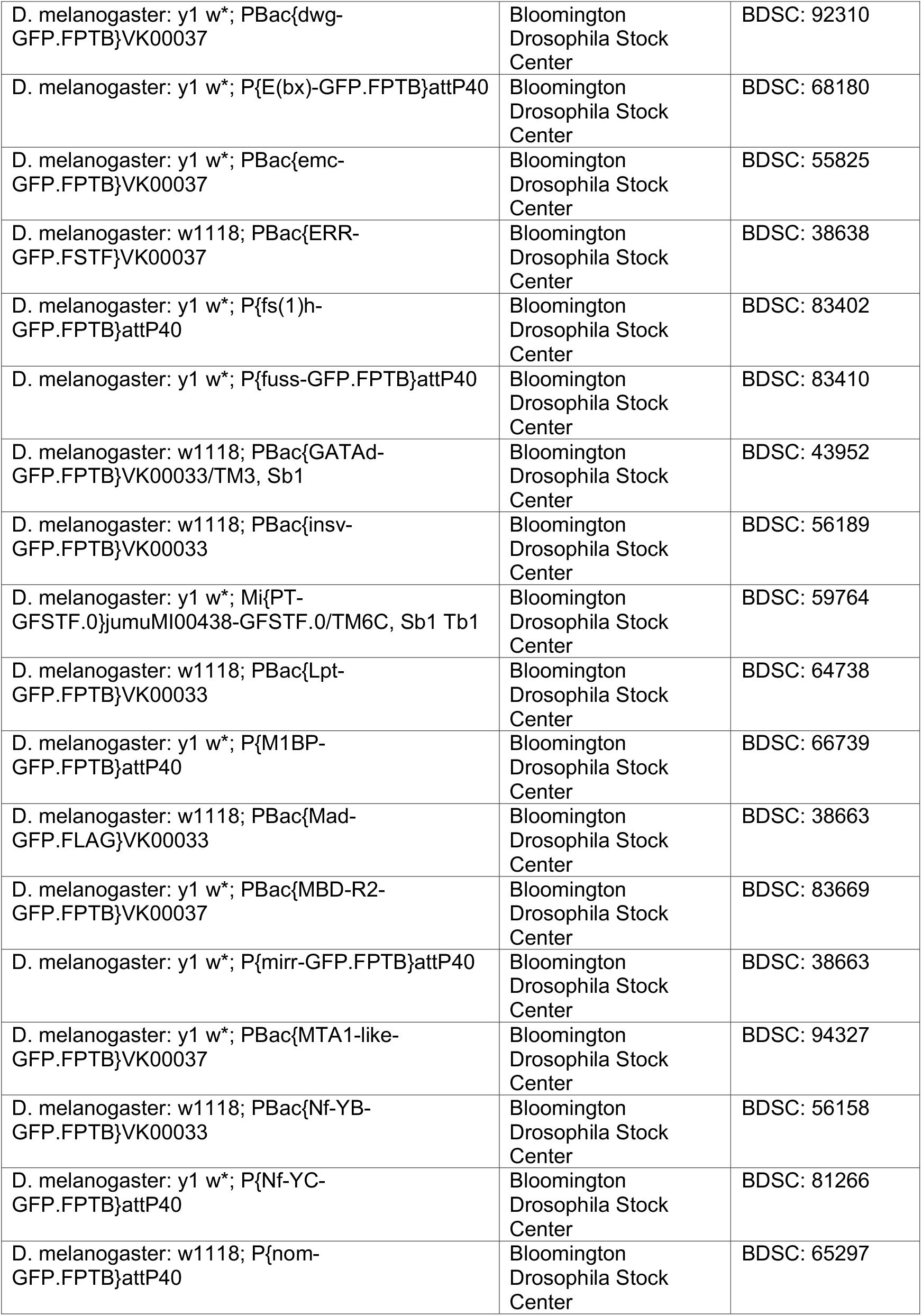
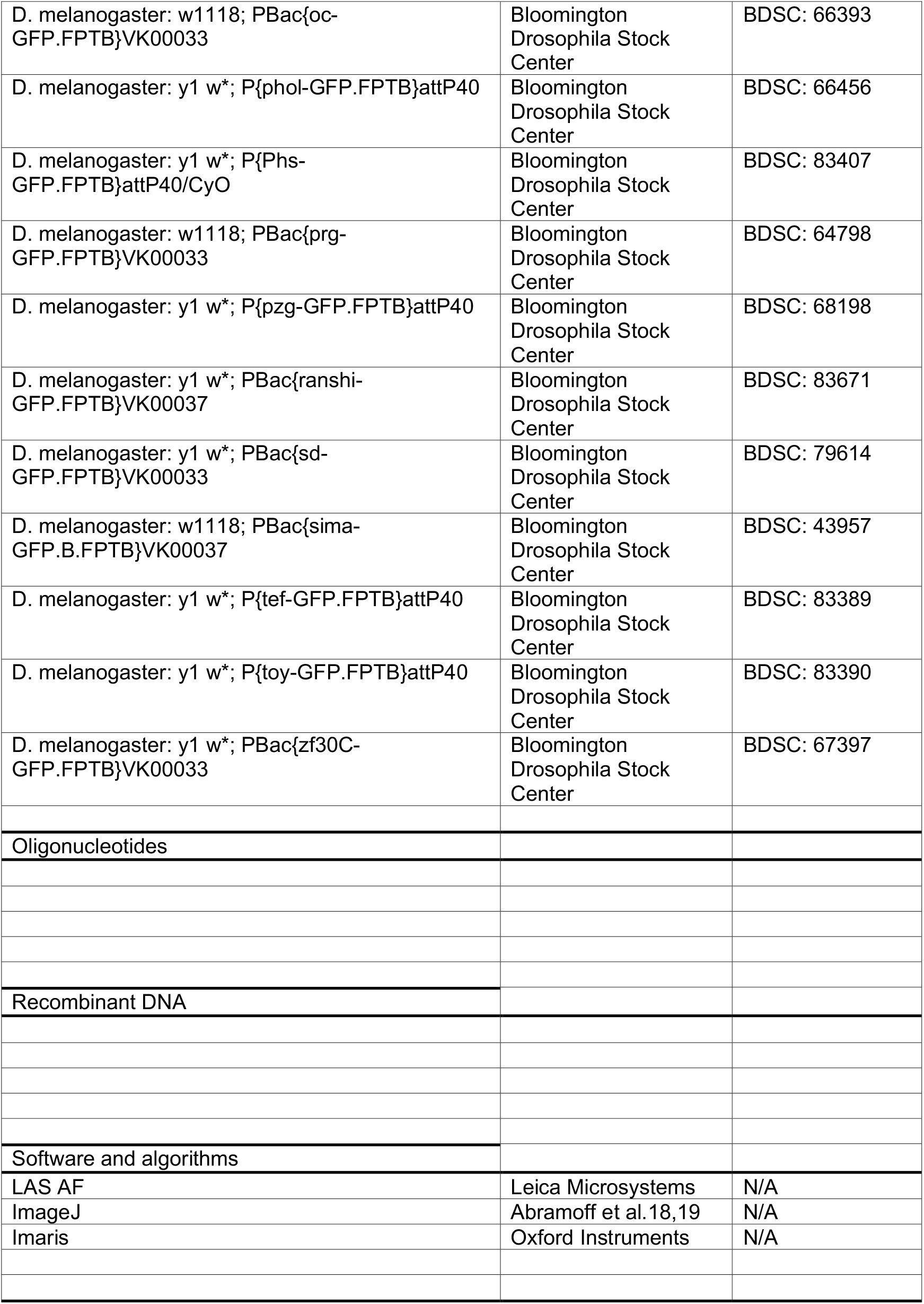
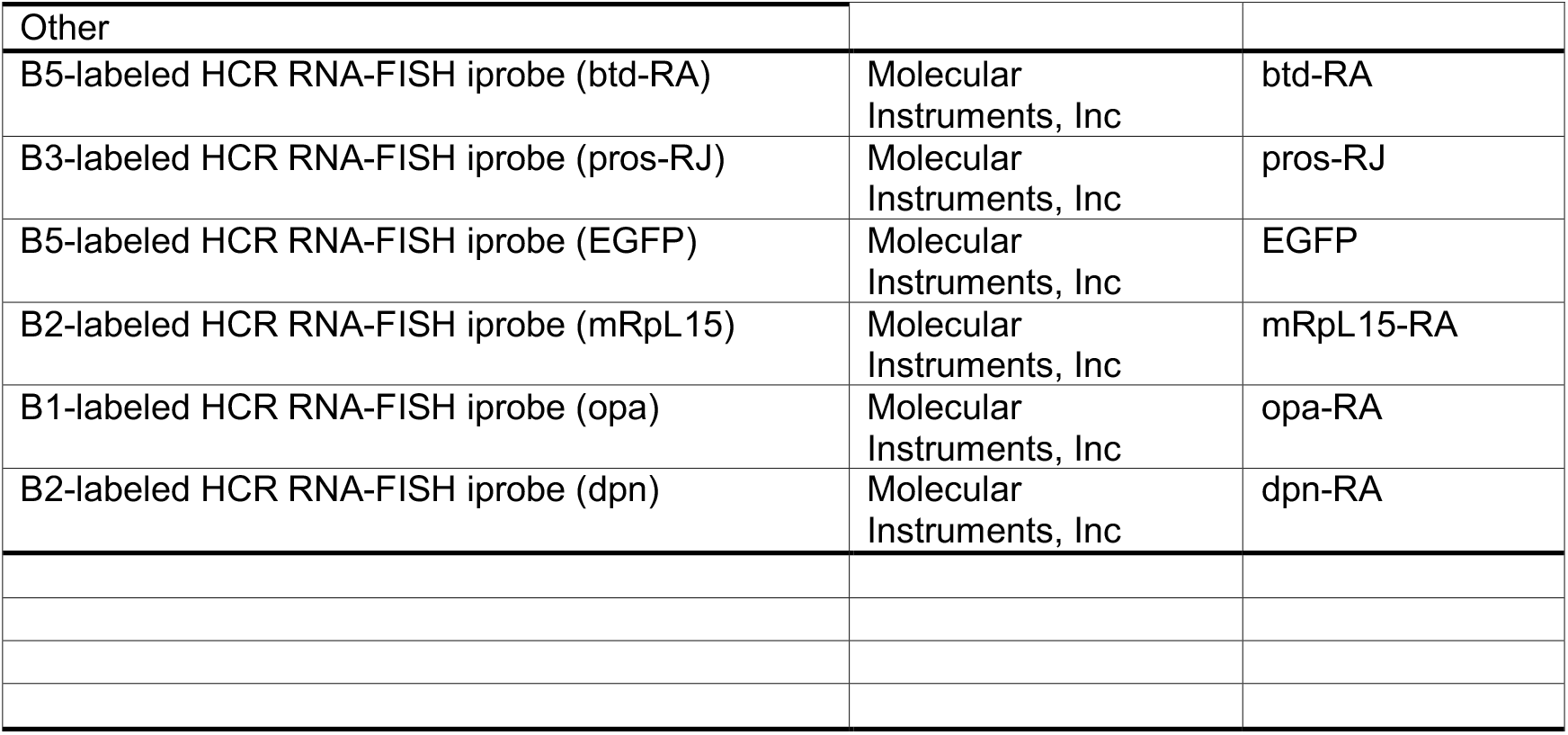

